# m⁶A-dependent FMRP control of DGKκ translation underlies core Fragile X phenotypes

**DOI:** 10.64898/2026.01.20.700356

**Authors:** Oktay Cakil, Boglarka Zambo, Tatiana Maroilley, Nathalie Drouot, Anastasiya Petrova, Luc Negroni, Gergo Gogl, Amélie Piton, Hervé Moine

## Abstract

Fragile X syndrome (FXS), a leading inherited cause of intellectual developmental disorder and autism, results from loss of the RNA-binding protein FMRP. Loss of FMRP causes excessive neuronal protein synthesis contributing to widespread functional disturbances, yet the mechanisms linking FMRP to specific mRNA targets remain unclear.

We show that FMRP promotes translation of the brain mRNA DGKκ by binding m⁶A-modified repetitive RNA motifs, thereby relieving a translational block encoded within its sequence. FMRP loss sharply reduces neuronal DGKκ, and DGKκ depletion alone reproduces hallmark FXS phenotypes, including hyperactivity, compulsive behavior, overgrowth, dendritic spine abnormalities, overactivated diacylglycerol signaling and increased protein synthesis. These findings identify DGKκ as a key effector of FMRP function and establish a regulatory axis where m⁶A RNA modification modulates neuronal translation. This work defines new principles of translational control in neurodevelopmental disorders and positions DGKκ as a central driver of FXS pathogenesis.

## Introduction

Fragile X syndrome (FXS) is one of the most frequent monogenic causes of intellectual disability (ID) and autism spectrum disorders (ASD). FXS has a prevalence of about 1:5000 males and 1:8000 females and is associated with comorbidities including hyperactivity, anxiety, hypersensitivity, stereotypies, epilepsy, connective tissue dysplasia, macro-orchidism, post-natal overgrowth and post-puberty facial dysmorphism ^1^. FXS is caused by the loss of expression or function of the *FMR1* gene encoding FMRP, an RNA binding protein implicated in very diverse biological functions such as chromatin remodeling, ion channel modulation, RNA stability, ribosome heterogeneity and translation regulation ^2–10^.

The *Fmr1*-KO mouse ^11^ recapitulates many FXS-like phenotypes (hyperactivity, altered anxiety, hypersensitivity, stereotypies, epilepsy, macro-orchidism, overgrowth, craniofacial anomalies) ^12^ associated to abnormal dendritic spine morphology across brain regions ^13–17^.

Global neuronal protein synthesis measurements show a net elevation in *Fmr1*-KO neurons ^18–21^, suggesting that FMRP primarily acts as a translational repressor. Although translational activation has paradoxically been reported for certain FMRP targets ^22,23^, the phenotypic alterations of the *Fmr1*-KO model are largely attributed to elevated neuronal protein synthesis due to loss of FMRP’s translational control ^12^.

How FMRP selects its target mRNAs to achieve such widespread neuronal translation dysregulation effect remains incompletely understood. While studies suggested that FMRP associates with mRNAs on a transcriptome-wide scale via short sequence abundant motifs ^24,25^, other studies have identified specific mRNAs bound via more complex less frequent motifs ^22,26,27^, leaving FMRP’s mRNA selection principles and translatome-wide impact unclear.

The diacylglycerol kinase kappa (*Dgkκ*) mRNA was evidenced as an mRNA most co-immunoprecipitated with FMRP in a CLIP (cross-linking and immunoprecipitation) assay performed in primary murine cortical neurons ^23,28^. Together with the fact that purified FMRP binds *Dgkκ* mRNA with unprecedent affinity compared to other proposed targets, this suggested that DGKκ might be a prominent target of FMRP and thus an important player in FXS ^23^. DGKκ protein level was found severely reduced in *Fmr1*-KO mouse and FXS brain extracts while its RNA level was unaffected ^23,28^, indicating that FMRP is required for the translation of DGKκ, in apparent contradiction with its primarily described function as a translation repressor ^29^. DGKκ could play an important role in FXS as it is a modulator of diacylglycerol (DAG) and phosphatidic acid (PA), two second messenger lipids playing pivotal roles in protein synthesis control and spine morphology dynamics and involved several signaling pathways altered in FXS ^30,31^. In agreement with a role in neuronal DAG signaling, FMRP loss disrupts mGluRI-dependent DAG signaling ^23^. Moreover, treatment of young adult *Fmr1*-KO mice with adeno-associated viral (AAV) vectors expressing a modified FMRP-independent DGKκ transgene provided long-term correction of the cerebral DAG/PA unbalance and of the FXS-related core behavioral phenotypes ^28^. These data suggest DGKκ loss contributes centrally to FXS pathomechanisms. However, DGKκ’s exact neuronal roles and FMRP’s control mechanism remained unknown.

Here we identify a previously unrecognized mechanism whereby FMRP regulates DGKκ translation by binding to a unique repeated RNA consensus motif that encodes a hard-to-translate amino acid sequence. We show that m^6^A modifications recruit FMRP and its homologs FXRs to resolve ribosome collisions on *Dgkκ* mRNA. In the *Fmr1*-KO brain, we identify region-specific deficits in DGKκ translation and demonstrate that DGKκ loss recapitulates FXS-like phenotypes, including altered neuronal morphology, signaling, behaviors, and physiology, and excessive protein synthesis — a hallmark of the disorder. Moreover, rare missense *DGKk* variants identified in individuals with intellectual developmental disorder support a conserved role in human neurodevelopment. Altogether, these findings establish FMRP as a positive translational regulator of DGKκ, a master controller of neuronal protein synthesis, thereby resolving a central paradox in FXS pathophysiology.

## Results

### FMRP binds to RNA sequence that encodes EPAP repeats

*Dgkκ* mRNA is a primary target of FMRP in primary murine cortical neurons and its expression is severely altered in both *Fmr1*-KO mice and human FXS patients brains ^23,28^. Identifying FMRP binding site on its RNA targets could help elucidate the mechanism(s) by which FMRP modulates translation.

DGKκ exhibits low expression levels, which poses challenges for conventional CLIP-seq methods to achieve adequate coverage of this gene. To address this issue, we adopted a HITS-CLIP strategy using a validated FMRP antibody (**Fig. S1A**) in hypothalamic neurons, where *Dgkκ* transcripts are 40-fold more abundant than in cortical neurons (**Fig. S1B**), to enhance sequencing coverage of *Dgkκ* mRNA. The robustness of the HITS-CLIP approach was supported by the detection of a single, well-defined FMRP-binding peak in the GC-rich exon 14 of *Fmr1* (**Fig. S1C**), the initial G-quadruplex–forming region with high affinity for FMRP ^27^, by the lack of enrichment of housekeeping genes like *Rplp0* mRNA (**Fig. S1C**), and by a significant overlap with previous FMRP-CLIP studies (**Fig. S1D**). Strikingly, a single strong FMRP peak was identified in the *Dgkκ* transcript, located within the coding sequence of exon 1 (**Fig. 1A**). This result was further confirmed using a modified iCLIP method (**Fig. S1E**). The FMRP binding site on *Dgkκ* mRNA spans a *∼*500-nucleotide region within exon 1, characterized by a high GC content and corresponds to more than 20 conserved repeats, preserved at both the RNA and protein levels, located downstream of a proline-rich region. The repetitive sequences are composed of a conserved GARCCGGCCCCA dodecamer motif (**Fig. 1A**) that encodes EPAP/EPDP amino acid repeats (this RNA sequence is thereafter called “EPAP”) (**Fig. S1F**). A reporter mRNA fused to exon 1 of *Dgkκ* was strongly bound by endogenous FMRP in Hela cells compared with a control construct lacking it, while deletion of the EPAP repeats [nt 316-874] abolished the interaction (**Fig. 1B**), confirming that the interaction of FMRP with this sequence is necessary.

**Figure 1:**
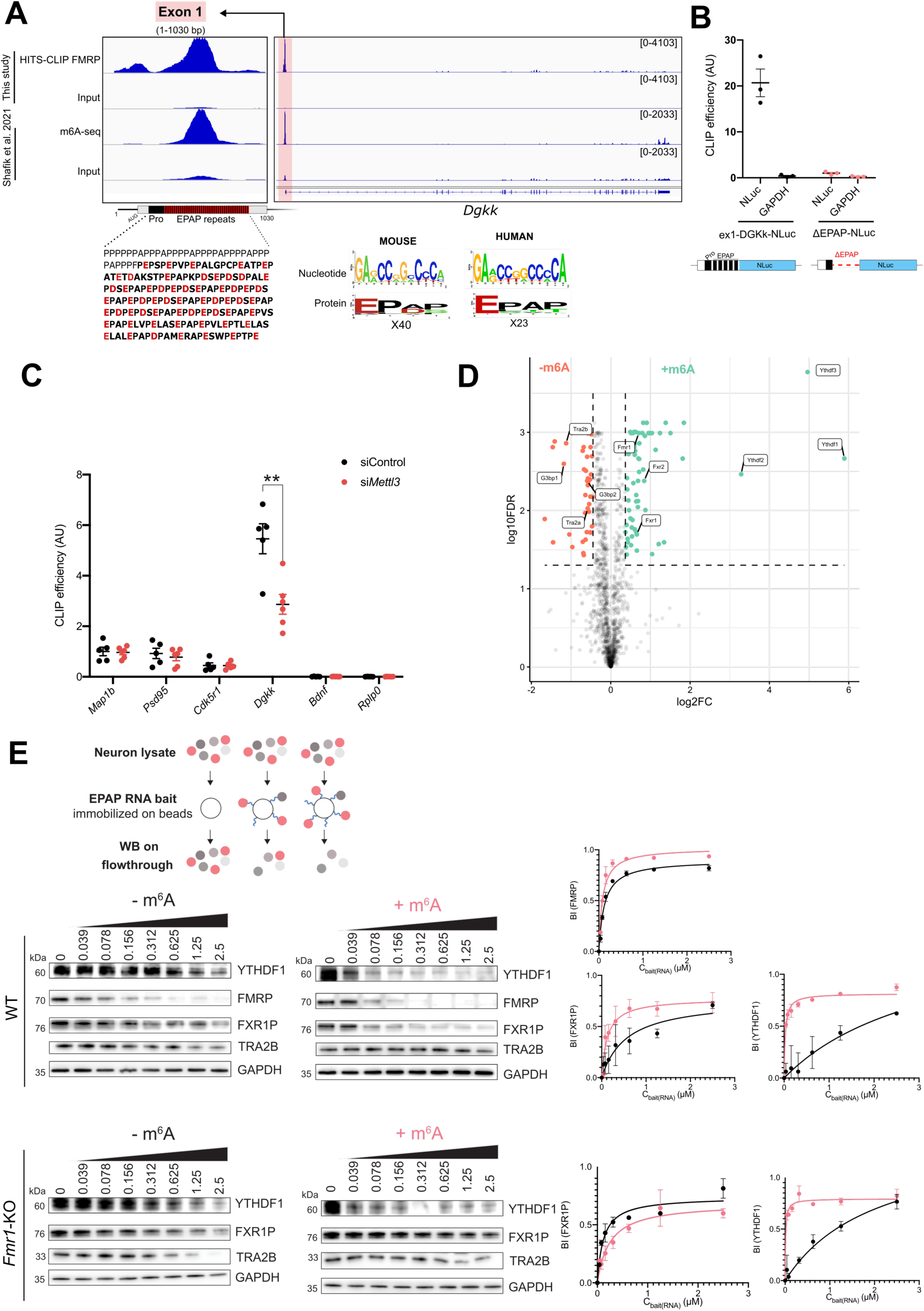
FMRP binds the m^6^A modified EPAP motif of *Dgkκ* mRNA. A) IGV representation of *Dgkκ* mRNA showing HITS-CLIP-seq profiles of FMRP (upper tracks) and m⁶A-seq profiles (lower tracks) from hypothalamic neurons, relative to input ^34^. Sequence logo analysis of conserved FMRP binding sites in mouse and human DGKκ. B) CLIP-qPCR in HeLa cells for Ex1-DGKκ-Nluc and ΔEPAP-Nluc constructs. Bars represent IP/Input relative enrichment. A non-target control (GAPDH) was included to assess specificity. Each dot represents an independent culture. C) CLIP–qPCR analysis of FMRP in cortical neurons treated with Acell siRNA control or siRNA targeting *Mettl3*. Bars represent IP/Input enrichment relative to Map1b signal. Well-established FMRP targets (*Map1b*, *Psd95*), non-target controls (*Rplp0*), an m^6^A-containing FMRP target (*Cdk5r1*), and a non-target (*Bdnf*) were included for specificity assessment. Each dot represents an independent culture. Data are mean ± SEM and analyzed using multiple t-tests comparing enrichment in WT and *Fmr1*-KO for each gene. **P < 0.01. D) RNA pull-down proteomic analysis using immobilized EPAP RNA probes with or without m^6^A modification, incubated with WT neuronal lysate. P-values were calculated using a two-sided unpaired t test, and statistical thresholds for binding (dotted lines) were determined at 1 standard deviation and P-value < 0.05. E) Schematic of the RNA nHU assay used to assess binding of EPAP RNA probes incubated with WT or *Fmr1*-KO neuronal lysates. Probes were incubated with lysates at concentrations ranging from 0 to 2.5 µM. Binding index (BI) values were analyzed by non-linear regression fitting to bimolecular binding equations.

### FMRP binds to “EPAP” RNA sequence in an m6A-dependent manner

A growing number of studies suggest that mRNAs bound by FMRP are enriched in m^6^A modifications ^9,32,33^. Interestingly, by querying the RMBase database (http://rna.sysu.edu.cn/rmbase), and reanalyzing brain m6A-seq ^34^, we identified a single strong peak of m^6^A RNA modification on *Dgkκ* mRNA, located in exon 1, centered on the EPAP repeats (**Fig. 1A**). This localization contrasts with the typical m^6^A distribution, predominantly found around the stop codon or in long internal exons ^35^, which suggested an alternative function.

To determine whether m^6^A modifications in *Dgkκ* mRNA are required for FMRP binding, we performed CLIP-qRT-PCR experiments in primary cortical neurons where METTL3, the catalytic component of the METTL3/METTL14 m^6^A writer complex ^36^, was silenced using siRNAs. We confirmed a nearly 50% reduction in METTL3 expression after 72 hours in neurons (**Fig. S1G**) and we assessed its impact on several known FMRP mRNA targets containing m^6^A, including *Map1b, Psd95, Cdk5r1,* and *Dgkκ*. We included *Bdnf* (methylated) and *Rplp0* (unmethylated) mRNAs, which are not known FMRP targets, as controls. We observed a significant upregulation of *Psd95* and a trend toward upregulation of *Cdk5r1*and *Bdnf,* as well as for *Dgkκ,* in agreement with the role of m^6^A in mRNA decay for those known modified mRNAs ^37^ (**Fig. S1H**). We then assessed the interaction between FMRP and these mRNAs. As expected *Dgkκ* exhibited a strong CLIP efficiency, in contrast with all tested mRNAs, but strikingly, only *Dgkκ* was affected by METTL3 knockdown (**Fig. 1C**). Specifically, CLIP/Input enrichment decreased by approximately 50% in METTL3 knockdown conditions compared to controls, indicating that FMRP binding to *Dgkκ* mRNA is dependent on m^6^A modifications.

To investigate whether FMRP binding to the “EPAP” sequence is direct and dependent on m^6^A, we analyzed the interaction of purified human FMRP isoform 1, the most abundant isoform, with a 76-nt synthetic RNA sequence containing 7 EPAP repeats bearing or not m^6^A. Using EMSA, we confirmed a direct interaction between FMRP and the (EPAP)7 RNA, with a twofold higher affinity for the methylated compared to the unmethylated form (**Fig. S1I**). These findings indicate that FMRP can bind the “EPAP” RNA motif *in vitro* and that its affinity is enhanced by m^6^A modifications.

m^6^A modifications recruit proteins that regulate RNA fate ^33,38–41^. Given that these proteins may influence DGKκ translational regulation, we conducted RNA-pulldown proteomic analysis using the (EPAP)7 RNA, with and without m^6^As, immobilized on streptavidin beads and incubated with cortical neuronal lysate. Interestingly, we observed enrichment of FMRP, as well as its paralogs FXR1P and FXR2P, with the methylated RNA (**Fig. 1D and supplementary Table S1**), consistent with the EMSA results. Additionally, we observed significant enrichment of the m^6^A readers YTHDF1, YTHDF3, and to a lesser extent, YTHDF2 (**Fig. 1D**). Moreover, the stress granule factor G3BP1 exhibited a preference for the unmethylated RNA, as previously demonstrated ^39^ (**Fig. 1D**). Analysis of RIP-seq datasets from m^6^A readers YTHDF1, YTHDF2, and YTHDF3 in primary hippocampal neurons ^42^ corroborated our RNA-pulldown results, with YTHDF1 and YTHDF3 showing higher binding affinity for *Dgkκ* mRNA compared to YTHDF2 (**Fig. S1J**). We complemented these data using RNA native hold-up (RNA-nHU), an interactomic method adapted from ^43^ that enables assessment of the relative binding affinities of an RNA ligand with its protein interactors within cell extracts (**Fig. 1E**). The RNA-nHU experiment was performed using both methylated and unmethylated (EPAP)7 RNA baits at concentrations ranging from 0 to 2.5 µM, incubated with cortical neuronal lysate. First, we confirmed that YTHDF1 exhibits significant affinity for m^6^A-containing RNA, which is exclusively dependent on the presence of m^6^A (**Fig. 1E**). Consistent with our RNA pulldown proteomics data, we also observed that m^6^A modification repels TRA2B binding, resulting in a higher affinity for the unmethylated RNA (**Fig. 1E**). We also confirmed the interaction of FMRP with the unmethylated (EPAP)7 and observed enhanced affinity for the methylated form (**Fig. 1E**). A similar binding was observed with the FXR1P paralog (**Fig. 1E**). The known heterodimerization of FXRs, combined with these findings, suggests that FXRs bind cooperatively to the m^6^A-EPAP motif. We therefore used RNA-nHU to investigate whether the absence of FMRP affects FXR1P and FXR2P binding. In *Fmr1*-KO cortical neurons, the interaction between FXR1P and the m^6^A-containing (EPAP)7 probe was decreased, suggesting that FXR1P, requires FMRP for a robust interaction with the “EPAP” (**Fig. 1E**).

### FMRP does not regulate *Dgkκ* mRNA transport and nuclear export

FMRP has been described to modulate neuritic transport and nuclear export of mRNAs ^32,44,45^. Since m^6^A was shown to influence mRNA neuritic transport and FMRP described as an m^6^A reader ^9,32,33^, we assessed whether the loss of FMRP could alter *Dgkκ* mRNA transport into neurites. For this purpose, we used primary cortical neurons grown on filters to physically separate soma from neurites and quantify mRNA levels by qRT-PCR in both fractions. The enrichment of *Col3a1* mRNA in neurites, a previously established transported mRNA, and the enrichment of *Gng3* mRNA in soma, known to be poorly transported, confirmed the quality of preparations (**Fig. S1K**). Our data showed that approximately 50% of *Dgkκ* mRNA is present in neurites, slightly exceeding the level observed for *Psd95* mRNA (∼40%). The absence of transport defect in *Fmr1*-KO neurons, indicated that *Dgkκ* mRNA transport into neurites is independent of FMRP (**Fig. S1K**).

Then, the nuclear export was assessed by isolating nuclei from the cytosol in WT and *Fmr1*-KO brain samples to quantify mRNA level by qRT-PCR on each fraction. The enrichment of nuclear U2 snRNA in nuclear fraction confirmed the correct separation of nuclei from cytosol. However, we detected no difference in *Dgkκ* mRNA levels between the nuclear and cytosolic fractions, indicating that FMRP does not influence the nuclear export of *Dgkκ* mRNA (**Fig. S1L**). These results suggest that FMRP is not involved in the regulation of *Dgkκ* mRNA neurite transport or nuclear export.

### EPAP repeats induce ribosome stalling

To determine the influence of “EPAP” sequence on DGKκ expression, we first assessed the consequence of the deletion of the exon 1 in *Dgkκ* mRNA (Δex1-DGKκ). Interestingly, deletion of this region resulted in a marked increase in DGKκ protein expression in Hela cells (**Fig. 2A**). Consistently, reporter NLuc construct containing this region (ex1-DGKκ-NLuc) displayed approximately sixfold lower expression than control construct, indicating that this region contains repressive element for DGKκ expression (**Fig. 2B**). “EPAP” RNA sequence encodes *∼*80% of prolines and acidic amino acids (aspartate D and glutamate E) (**Fig. 1A**), well-established to cause ribosome stalling ^46–49^, suggesting a ribosomal stalling could be imposed by the protein sequence itself. To test this hypothesis, and to distinguish the contributions of the RNA sequence itself from its role in protein coding, we introduced reading-frame shifts that altered the resulting polypeptide sequence while preserving the overall RNA sequence. As expected, disrupting the proline-rich motif (fs_-1_Pro_+1_-NLuc) upstream of the EPAP repeats increased NLuc activity, consistent with a relief of peptide-mediated translational repression (**Fig. 2B**). Strikingly, frame shifts at the EPAP repeats (fs_-1_EPAP_+1_-NLuc, fs_+1_EPAP_-1_-NLuc) produced a comparable increase in NLuc activity, indicating that both proline-rich and EPAP repeats exert an inhibitory effect primarily through their decoding rather than at their RNA sequence level (**Fig. 2B**). Because impaired translation elongation promotes ribosomal collisions ^50^, we next investigated whether the “EPAP” promotes such collisions. To this end, we performed RNase protection assays to detect ribosome-protected RNA fragments within the *Dgkκ* exon 1 mRNA sequence. Treatment of cells with a low concentration of the elongation inhibitor emetine, which enhances ribosomal collisions ^51^, increased the abundance of disomes in RNase A–digested lysates fractionated on a sucrose gradient. (**Fig. 2C**). Accordingly, using a radiolabeled probe targeting the luciferase sequence, we detected disome footprints of ∼60 nt upon emetine treatment, in contrast to the ∼30 nt monosome footprints observed in control condition (**Fig. 2C**). Strikingly, fusion of the exon 1 of DGKκ upstream the NLuc reporter resulted in a strong depletion of monosome within the luciferase sequence, consistent with a strong translational roadblock (**Fig. 2D**). Moreover, when using an EPAP-specific probe, we detected ribosome footprints corresponding to higher-order collided ribosomes. Importantly, this signal was dependent on the EPAP coding frame: changing the reading frame within the EPAP motif markedly reduced the abundance of higher-order collision footprints relative to monosome footprints (**Fig. 2D**) and slightly increase the downstream monosome signal within the luciferase sequence, thereby excluding a contribution from RNA secondary structure. Together, these data demonstrate that the EPAP-encoding region of *Dgkκ* promotes ribosomal collisions in a reading frame–dependent manner, consistent with a mechanism of translation repression.

**Figure 2:**
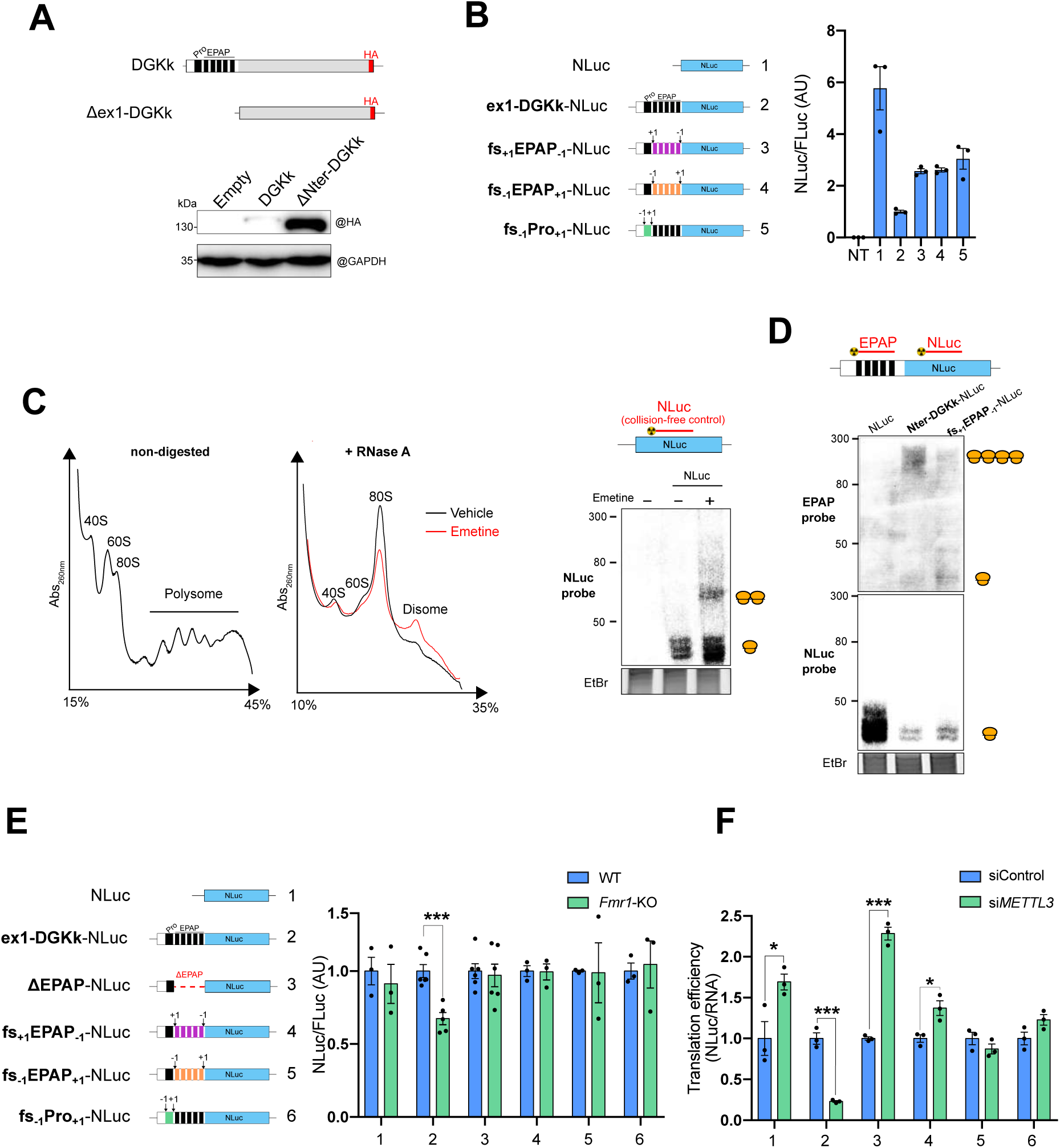
FMRP and m6A mitigates ribosomal stalling imposed by the EPAP repeats. A) Representative western blot of transfected WT DGKκ and ΔNter DGKκ tagged with HA in HeLa cells. GAPDH was used as a loading control. B) Translation efficiency measurements of the indicated constructs in HeLa cells. Luciferase signals were normalized to RNA levels of the corresponding constructs. C–D) Sucrose gradient control shows ribosome profile of undigested and RNase-A-digested lysate. Northern blot analysis of RNase-A–digested ribosomal footprints. Specific radio-labeled probes for NLuc (C-D) and EPAP (D) were used. For emetine-positive control, cells were incubated with 2 µM emetine for 15 min prior to lysis. Ethidium bromide staining was used as a loading control. E) Luciferase assays of indicated NLuc constructs in WT and *Fmr1*-KO cortical primary neurons at DIV4. Firefly luciferase was used as a normalization control. Data are mean ± SEM and analyzed using multiple t-tests comparing enrichment in WT and *Fmr1*-KO for each gene. ***P < 0.001. F) Translation efficiency measurement of indicated constructs in HeLa cells transfected with control or *METTL3*-targeting siRNA. Luciferase signals were normalized to RNA levels of the corresponding constructs. Data are mean ± SEM and analyzed using multiple t-tests comparing enrichment in WT and *Fmr1*-KO for each gene. *P<0.05; ***P < 0.001.

### FMRP alleviates ribosomal stalling imposed by EPAP repeats

To better characterize the mode of action of FMRP in controlling *Dgkκ* mRNA translation, we overexpressed an ex1–DGKκ–NLuc reporter in wild-type and *Fmr1*-KO neurons. Remarkably, we partially recapitulated FMRP-dependent translational control, observing ∼40% reduced luciferase activity in *Fmr1*-KO neurons (**Fig. 2E**). Deleting the EPAP repeats abolished this regulation. Likewise, frameshift mutations in the EPAP repeats (−1/+1 and +1/−1), which preserve the RNA sequence but alter the encoded protein, eliminated FMRP-dependent control (**Fig. 2E**). Changing the reading frame within the proline-rich sequence similarly disrupted regulation, demonstrating that the encoded protein sequence–rather than the RNA sequence itself–is essential for FMRP-mediated translational enhancement (**Fig. 2E**).

Together, these data reveal that FMRP promotes context-dependent translation at sequences encoding repetitive, sub-optimal amino acids.

### m^6^A enhances translation efficiency within the EPAP repeat context

The conserved m⁶A consensus motif within the EPAP domain in both mouse *Dgkκ* and human *DGKκ* corresponds to the GA**A**CCN context encoding the E-P dipeptide, with methylation occurring on the GA**A** codon. Recent studies indicate that m⁶A modifications within coding sequences, particularly on GA**A** codon, slow translation elongation and ultimately trigger ribosomal collisions ^52–54^.

To investigate the impact of EPAP repeats on DGKκ expression, we used the nanoluciferase gene reporter system described above in non-neuronal cells. Cells were treated with *METTL3* siRNA which resulted in a ∼2.5-fold increase in ex1-DGKκ-NLuc RNA levels, markedly higher than that observed for the control NLuc construct (**Fig. S2A**). Notably, this increased RNA level was specific to the EPAP repeats, as its deletion in the ΔEPAP-NLuc construct, abolished this effect (**Fig. S2A**). These results are consistent with previous studies showing that m^6^A modifications within the coding sequence promote efficient mRNA degradation ^55^.

Surprisingly, however, luciferase activity from the ex1-DGKκ-NLuc construct did not correlate with RNA abundance, leading to an ∼80% reduction in translation efficiency (luciferase/RNA ratio) when silencing METTL3 compared with the control condition (**Fig. 2F**). This effect was also seen, in a dose-dependent manner using STM2457 METTL3 specific inhibitor (**Fig. S2B**). Interestingly, this effect that was not observed when the EPAP repeats or the proline-rich region were out of frame (**Fig. 2F**) and led to an increase or no change of translation efficiency in accordance with previous reports ^53,54,56,57^ .

Moreover, this effect is independent of YTHDF1 and YTHDF2, as their silencing increased RNA level of ex1-DGKκ-NLuc construct, without affecting its translation efficiency, consistent with their well-established roles in regulating mRNA stability ^35^ (**Fig. S2C**). Overall, these results indicate that, within the EPAP repeat context, m^6^A modifications counteract a sequence-imposed translational repression and positively modulate translation efficiency.

### DGKκ deficit varies across brain despite conserved *Dgkκ* mRNA/FMRP interaction

Our prior work showed that FMRP loss strongly impairs DGKk translation in *Fmr1*-KO cortical neurons ^23,28^. Given DGKk’s heterogeneous brain expression (**Fig. 3A**), neuron-specificity (**Fig. 3B**), and binding by all FXR paralogs to the EPAP repeats (**Fig. 1D**), we investigated whether the expression of DGKk was differentially altered across distinct brain regions.

**Figure 3:**
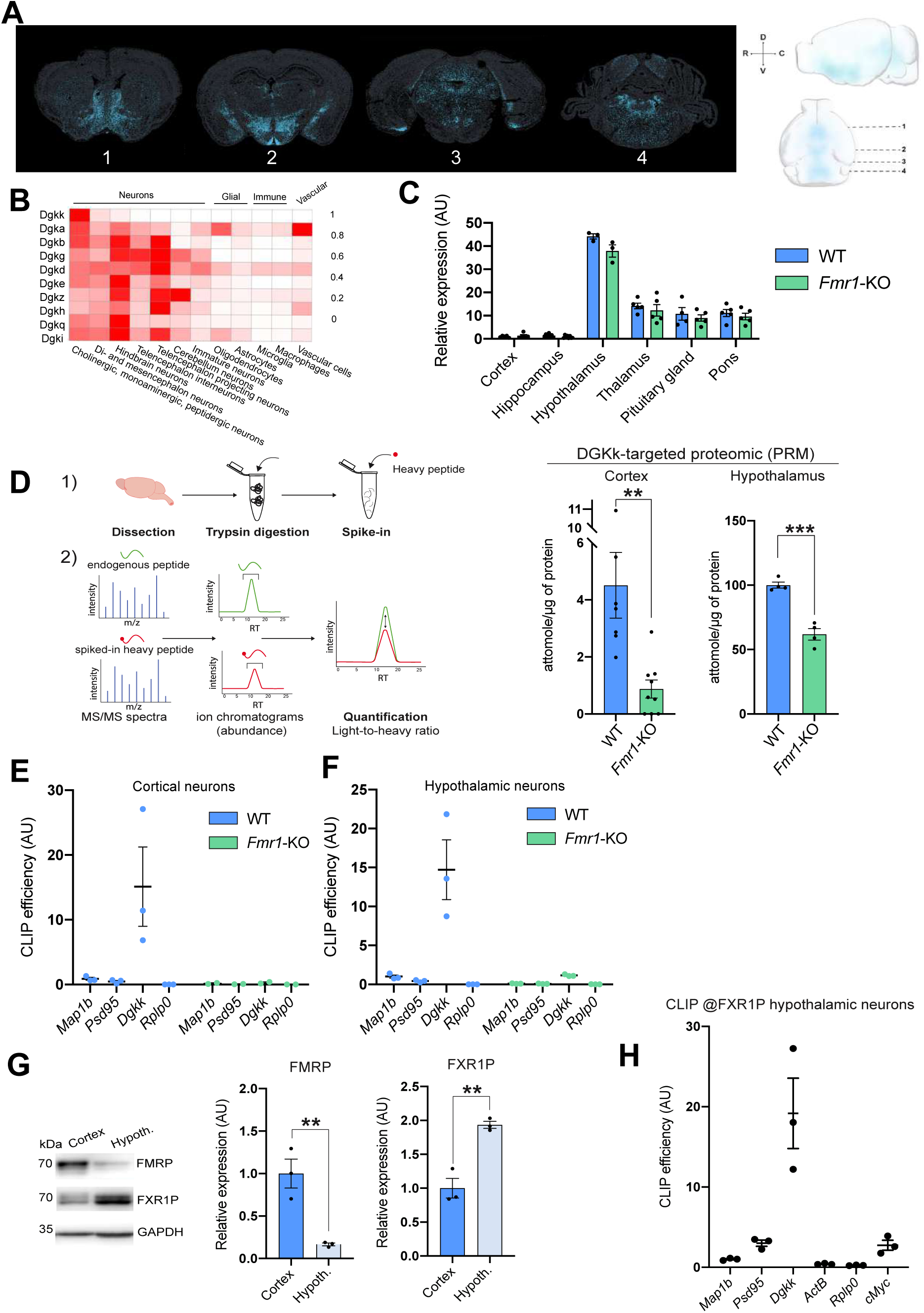
FXRs bind *Dgkκ* mRNA across the brain and display redundant function to regulate its translation. A) Single-molecule–resolution spatial transcriptomic MERFISH data (Vizgen) showing *Dgkκ* mRNA (blue) in mouse brain coronal sections. Upper panel: 3D reconstruction generated from 67 MERFISH sections. B) Single-cell transcriptomic ^137^ reanalysis showing the expression patterns of DGKs across major brain cell types. C) qRT–PCR analysis of *Dgkκ* expression in cortex, hippocampus, hypothalamus, thalamus, pituitary gland, and pons from adult WT and *Fmr1*-KO mice. Fold changes were calculated using the ΔΔCt method with Tbp as the housekeeping gene. Data are mean ± SEM and were analyzed using multiple t-tests comparing expression in WT and *Fmr1*-KO across brain regions. D) PRM-targeted proteomic quantification of DGKκ in cortex and hypothalamus from WT and *Fmr1*-KO mice. Each dot represents an individual mouse. Data are mean ± SEM and were analyzed using an unpaired Student’s t-test. **P < 0.01, ***P<0.001. E–F) CLIP–qPCR analysis of FMRP in primary E) cortical and F) hypothalamic neurons at DIV8. Bar graphs represent IP/Input enrichment relative to the Map1b signal. Established FMRP mRNA targets (*Map1b* and *Psd95*) and a non-target control (*Rplp0*) were included to assess specificity. Analysis in *Fmr1*-KO neurons confirms immunoprecipitation specificity. Each dot represents an independent culture. G) Representative western blots of FMRP and FXR1P in cortex and hypothalamus. Quantification was normalized to GAPDH and expressed relative to cortex expression. Data are mean ± SEM and were analyzed using unpaired Student’s t-tests. **P < 0.01. H) CLIP–qPCR analysis of FXR1P in primary hypothalamic neurons at DIV8. Bar graphs represent IP/Input enrichment relative to the Map1b signal. An established FXR1P target (*cMyc*) was used as a positive control, while Rplp0 and Actb served as non-target controls.

Detection of DGKκ by conventional methods is challenging due to its low protein abundance and the absence of specific commercial antibodies. To assess the consequence of FMRP loss on DGKκ protein level, we used targeted mass spectrometry called Parallel Reaction Monitoring combined with stable isotope-labeled standard peptides (PRM-SIS) to assess DGKκ absolute level. While there was no difference in *Dgkκ* mRNA level between *Fmr1*-KO and WT across the brain (**Fig. 3C**), DGKκ protein was reduced by ∼80% in *Fmr1*-KO cortex and primary cortical neurons, contrasting with a ∼50% decrease in the hypothalamus (where DGKκ is most highly expressed) (**Fig. 3D, Fig. S3A, B**). This observation prompted us to determine whether the interaction between FMRP and *Dgkκ* mRNA was conserved across different brain regions. Thus, using a CLIP approach to compare FMRP interaction in cortical neurons, where DGKκ expression is very low, to hypothalamic neurons, where *Dgkκ* is over 40-fold higher (**Fig. 3C**). Surprisingly, CLIP-qRT-PCR analyses revealed a strong and similar enrichment of *Dgkκ* mRNA relative to input similar in both neuron subtypes (**Fig. 3E**, **F**). This suggests that *Dgkκ* mRNA is a favored target of FMRP compared with other proposed targets such as *Map1b* and *Psd95,* across different brain regions and independently of its mRNA level.

The FXR paralogs (FMRP, FXR1, FXR2) share ∼65% sequence homology across mammals, including conserved KH1, KH2, and RGG RNA-binding domains ^58^, suggesting functional redundancy. *Fmr1* mRNA level was found ∼4-fold higher in cortex than hypothalamus, whereas *Fxr1* showed the inverse pattern (**Fig. S3C**), suggesting that FXRs may display reciprocal expression across brain regions. These differences persisted at the protein level, with opposing FMRP/FXR1 abundance profiles (**Fig. 3G**). To test whether FXR1P binds and regulates *Dgkκ* mRNA we performed CLIP-FXR1P in primary hypothalamic neurons where *Dgkκ* expression peaks. *cMyc* mRNA – a known FXR1P target ^59^ – was significantly enriched relative to input, IgG, and negative control *Rplp0*. *Dgkκ* mRNA showed 5- to 10-fold greater enrichment than all tested targets, including *Map1b*, *cMyc*, and *Psd95* (**Fig. 3H**), confirming the specific and strong binding of FXR1P to *Dgkκ* mRNA. These data suggest a compensatory mechanism by FXR paralogs in DGKκ-rich regions that may account for the partial loss of DGKκ protein levels in *Fmr1*-KO mice.

### *Dgkκ*-KO mouse recapitulates core FXS behavioral phenotypes

To assess the biological role of DGKκ, we generated a *Dgkκ* knockout mouse (*Dgkκ*-KO) via CRISPR-Cas9 deletion of exon 8, inducing a frameshift and premature stop in exon 9 (**Fig. S4A-D**). No transcriptional compensation occurred among the nine other DGK isoforms (**Fig. S4E**). *Dgkκ*-KO mice appeared grossly normal, with no reproductive bias or congenital malformation–including absence of hypospadias, in contrast to previous proposal based on GWAS ^60^. *Dgkκ*-KO mice exhibited progressive body weight gain from 3-22 weeks (**Fig. 4A**), driven by lean mass (**Fig. 4B**) and fluid increases (**Fig. 4C**)—not fat (**Fig. 4D**) or size (qNMR; **Fig. 4E**)—plus elevated bone density (**Fig. 4F**), heart weight (**Fig. 4GH**), and compensated cardiomyopathy (**Fig. S4FG**). Unlike *Fmr1*-KO mice and FXS patients, macroorchidism was absent ^11,61^. Thus, DGKκ loss recapitulates syndromic FXS-like overgrowth ^62^, while other FMRP targets likely mediate macroorchidism.

**Figure 4:**
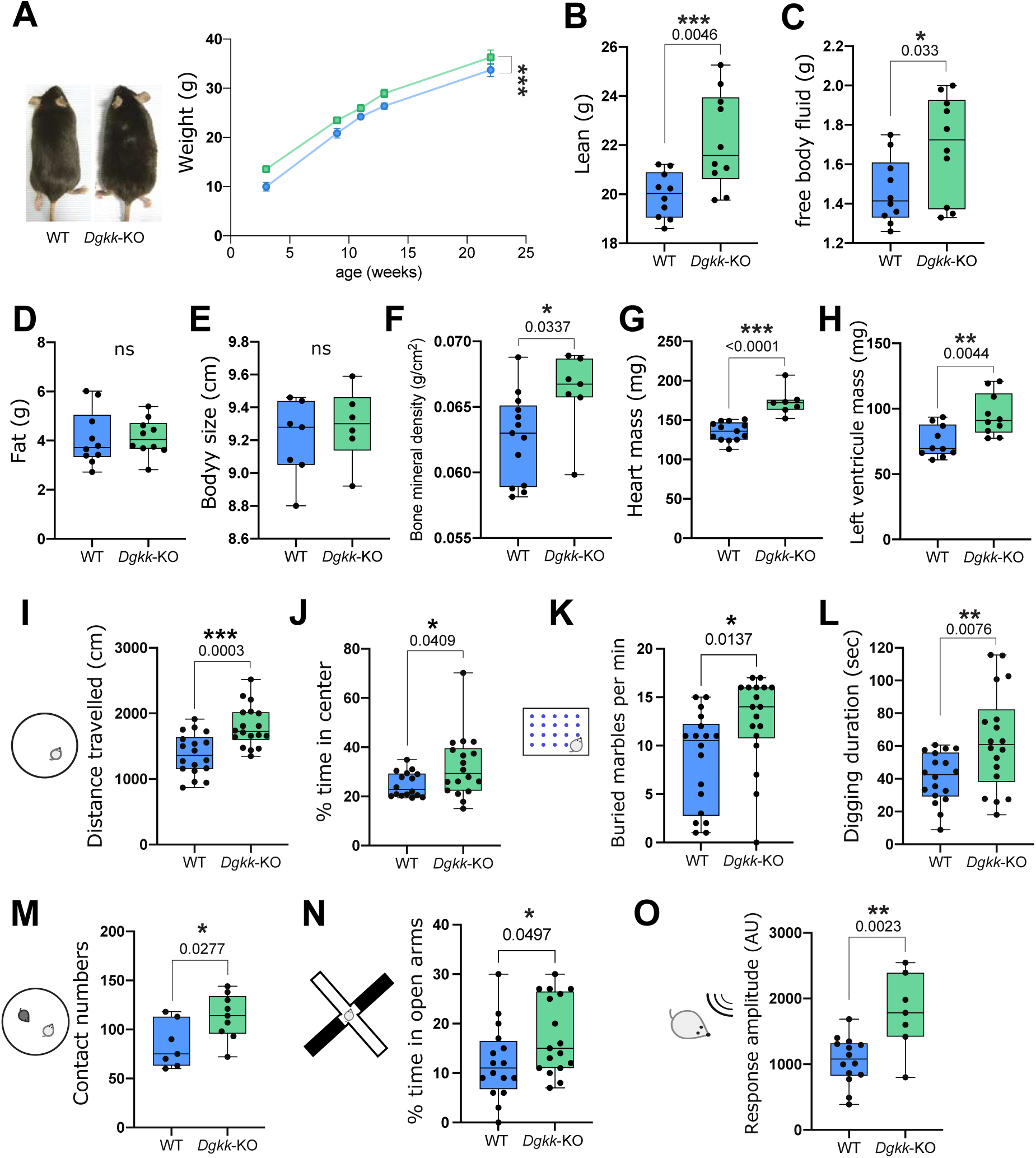
D*g*kκ-KO mice exhibit FXS-like and overgrowth phenotypes. A) Body weight of WT and *Dgkκ*-KO mice measured across multiple ages, with representative images of WT and *Dgkκ*-KO mice (scale bar, 1 cm). Data are presented as median with interquartile range with minimum and maximum values and analyzed by two-way repeated-measures ANOVA to evaluate the overall effect of genotype. ***P < 0.001. B-F) Quantitative NMR (qNMR) body composition analysis showing B) lean mass, C) free body fluid mass D), fat mass, E) body size measurements, F) bone density. G, H) Echocardiographic assessment of G) heart mass and H) left ventricular mass. Each dot represents one mouse. Data are medians with interquartile range with minimum and maximum values and were analyzed using unpaired Student’s t-tests. **P<0.01; ***P<0.001. I-J) Open field test. I, total distance traveled over 30 min and J, percentage of time spent in the center of the arena. K) Marble burying test. Number of marbles buried within 15 min, indicative of compulsive-like behavior and/or hyperactivity. Data are presented as median with interquartile range with minimum and maximum values and analyzed by unpaired Student’s t-tests. *P<0.05. L) Digging behavior measurement. M) Social interaction test. Frequency of interactions between the test mouse and a novel conspecific. N) Elevated plus maze test. Percentage of time spent in open arms, reflecting anxiety-like behavior. O) Acoustic startle response. Amplitude of the startle reflex elicited by an acoustic stimulus. B-0) Each dot represents one mouse. Data are represented as median with interquartile range with minimum and maximum values and were analyzed using unpaired Student’s t-tests, *P<0.05, **P<0.001, ***P < 0.001.

At 13 weeks, *Dgkκ*-KO male mice underwent a battery of behavioral and cognitive tests. Use of open field test revealed an increased locomotor activity of *Dgkκ*-KO compared to WT mice (**Fig. 4I**) with increased time spent in the center of the arena (**Fig. 4J**) that were evocative of a hyperactivity behavior with reduced anxiety. Hyperactivity was also supported by the increased marble burying (**Fig. 4K**). Increased digging duration and number of diggings suggested compulsive-like behavior (**Fig. 4L**). An increased contact numbers and duration in the social interaction test suggested increased sociability (**Fig. 4M, Fig. S4H**), but could also resulted from hyperactivity and reduced anxiety. Increased time in the open arm in the elevated plus maze with increased arm entries trends suggested also hyperactivity and reduced anxiety (**Fig. 4N, Fig. S4I**). A trend for alteration of recognition ability was observed in the novel object recognition test (**Fig. S4J**), suggesting *Dgkκ*-KO mice do not present major episodic-like memory alteration. No nesting behavior or social memory defects were detected (**Fig. S4KL).** Finally, the startle response amplitude after pre-pulse Inhibition (PPI) was also found significantly increased (**Fig. 4O**), suggesting that *Dgkκ*-KO mice have a sensorimotor gating deficit and a hypersensitivity to acoustic stimuli. Collectively, these findings underscore the impact of DGKκ loss on multiple aspects of mouse cognitive and behavioral function, including hyperactivity, hypersensitivity, and compulsive-like behaviors, highlighting its role in neurodevelopment. Notably, these phenotypes overlap with those observed in the *Fmr1*-KO mouse model.

### DGKκ loss disrupts spine morphology, mGluRI-dependent activity, and leads to excessive neuronal translation

The neuron-specific expression of DGKκ (**Fig 3B**) and the neurodevelopmental behavioral alterations caused by its depletion suggested that DGKκ plays important neuronal functions. To identify the neuronal subcellular localization of DGKκ, we conducted an overexpression experiment in primary hippocampal neurons and performed immunostaining against HA-tagged DGKκ at DIV17. DGKκ was predominantly found enriched at the plasma membrane, similar to when it was overexpressed in non-neuronal cells ^63^. Within dendrites, DGKκ was found specifically enriched in post-synaptic densities of mature spines colocalizing with the synaptic marker PSD95 (**Fig. 5A**) and ribopuromycylation signal (**Fig. 5B and S5A**). This finding supports a role of DGKκ in synaptic function and suggests a contribution in modulating dendritic spines local translation.

**Figure 5:**
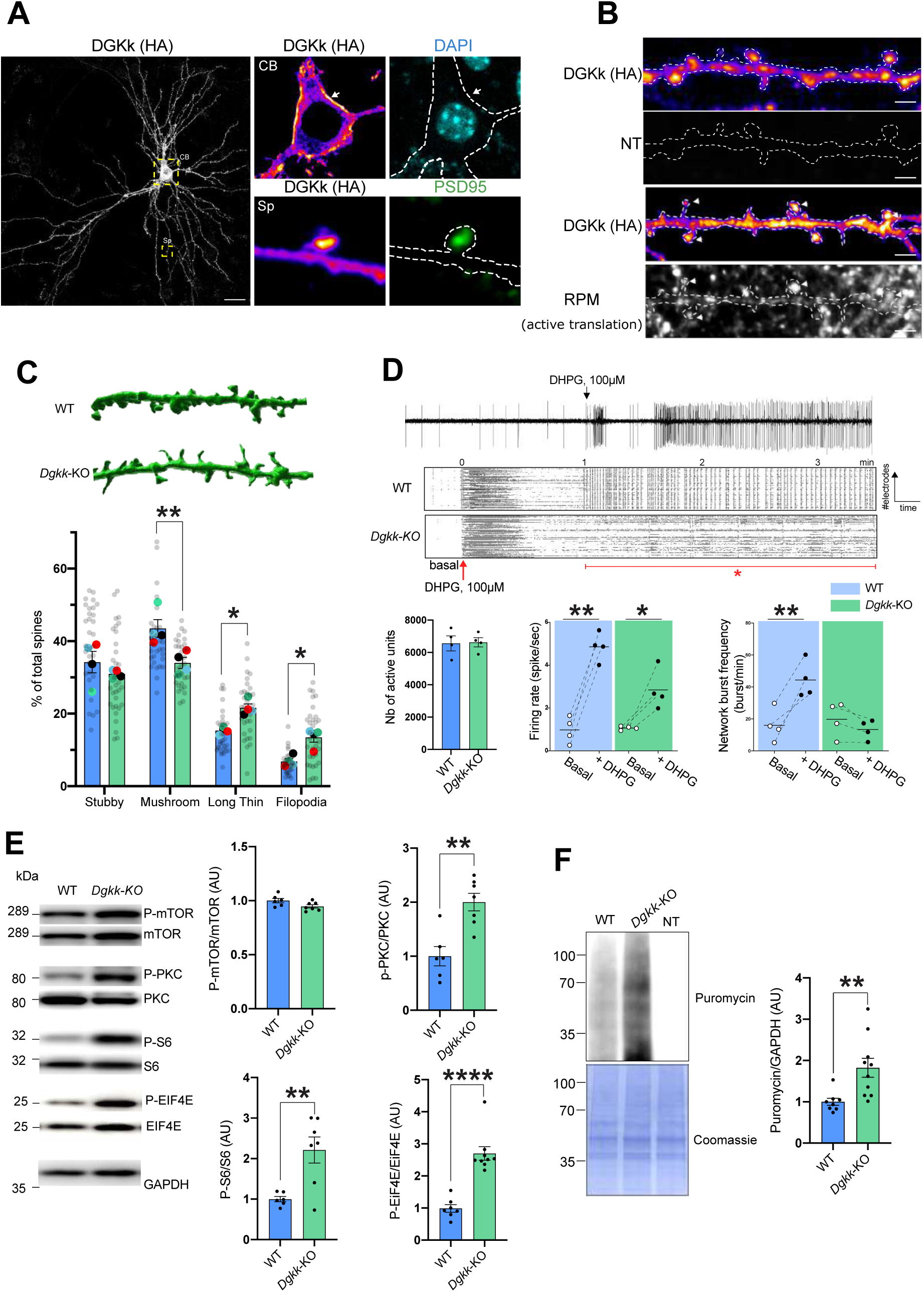
DGKκ loss results in dendritic spines morphology and mGluRI-dependent activity alterations, and aberrant DAG-signaling leading to increased neuronal translational rate. A) Neuronal localization of HA-DGKκ (fire) in primary hippocampal neurons and its colocalization with the synaptic marker PSD95 (green) at 17 DIV. Hippocampal neurons were magnetofected at 3 DIV with HA-tagged DGKκ constructs. At 17 DIV, neurons were immunolabeled with antibodies against HA and PSD95. Images were acquired by confocal microscopy. Digital magnification shows colocalization of DGKκ with PSD95 in dendritic spines. CB: cell body; Sp: spine. Scale bar: 30 µm. B) Ribopuromycylation assay in hippocampal neurons at DIV17. Top panel: no puromycin control. Bottom panel: arrows indicate colocalization of DGKκ (fire) with puromycin signal (white), highlighting sites of active translation. Scale bar: 2 µm. C) Quantification and classification of dendritic spine morphology in magnetofected GFP-positive hippocampal neurons at 21 DIV from WT and *Dgkκ*-KO mice. Upper panel: representative 3D reconstructions of dendritic spine morphology for each condition. Spine analysis was performed using Imaris on GFP-labeled neurons, analyzing 100–200 spines per 8–10 neurons per biological replicate. Each colored dot represents an individual culture. Data are presented as mean ± SEM and analyzed using two-way ANOVA followed by Sidak’s multiple-comparisons test. *P < 0.05; **P < 0.01. D) MEA recordings. Upper panel: raster plot example in which each line represents a different electrode (subset shown) and each dot represents an above-threshold spiking event. Lower panel: mGluRI-dependent activity in WT and *Dgkκ*-KO cortical neurons measured before and after 1 min of treatment with 100 µM DHPG. Each dot represents data from an individual embryo. Data are mean ± SEM and analyzed using paired Student’s t-test. *P < 0.05; **P < 0.01. E) Representative western blots of lysates from WT and *Dgkκ*-KO cortical neurons at 7 DIV and quantification of phosphorylation and total levels of indicated proteins. GAPDH was used as a loading control. Phospho-protein signals were normalized to total protein and expressed relative to WT. Data are mean ± SEM and were analyzed using unpaired Student’s t-tests. **P<0.01; ****P<0.0001. F) Representative western blots of puromycin-labeled neuron lysates from WT and *Dgkκ*-KO cortical neurons treated for 30 min with 20 µg/mL puromycin. Puromycin signals were normalized to GAPDH and expressed relative to WT. Each dot represents an individual culture. Data are mean ± SEM and were analyzed using unpaired Student’s t-tests. **P<0.01.

Dendritic spine morphology is frequently altered in neurodevelopmental disorders and these morphological alterations are proposed as an underlying cause of these disorders ^64^. To analyze the effects of DGKκ loss on dendritic spine morphology, primary hippocampal neurons from WT and *Dgkκ*-KO mice were magnetofected at 4DIV with a GFP-expressing plasmid and imaged at DIV21. A significant decrease in mature “stubby” and “mushroom” spines alongside a notable increase in immature “long thin” and “filopodia” spines was observed in *Dgkκ*-KO neurons (**Fig. 5C**).

Given the cognitive alterations of *Dgkκ*-KO mice, the increased proportion of immature spines in *Dgkκ*-KO neurons, and the consequences of other DGKs loss on synaptic plasticity ^65^, we hypothesized that the loss of DGKκ would likely lead to synaptic transmission dysfunction. To test this hypothesis, we recorded the neuronal activity of WT and *Dgkκ*-KO primary cortical neurons at DIV21 (time at which neural activity was highest **Fig. S5B**) on high-density multi-electrode array (HD-MEA) chips. The maturity of the culture was assessed by the spiking frequency of individual neurons, the burst neuronal network activity (**Fig. S5C**), and the responsiveness to (S)-3,5-Dihydroxyphenylglycine (DHPG), a specific agonist of group 1 metabotropic glutamate receptors (mGluRIs) (**Fig. 5D**).

While the number of active units was consistent between WT and *Dgkκ*-KO neurons, reflecting a comparable number of cells on the HD-MEA chip, at basal level, we observed similar firing rates and burst network frequencies between WT and *Dgkκ*-KO neurons, suggesting no apparent difference in absence of stimuli (**Fig. 5D**). However, upon neuronal activation with DHPG, we observed a 5-fold increase in neuronal firing rate for WT neurons, compared to only a 3-fold increase for *Dgkκ*-KO neurons. Furthermore, WT neurons showed an increased synchronization of their network activity whereas *Dgkκ*-KO neurons showed a loss of synchronization (**Fig. 5D**). Overall, the absence of DGKκ in neurons disrupts their intrinsic firing activity and synaptic transmission, resulting in alterations to the neuronal network coherence in vitro.

DGKs convert DAG into PA and are thought to play a key role in neuronal signaling by attenuating DAG-mediated pathways ^65^. To define the molecular alterations caused by the loss of DGKκ, we analyzed the activation status of several members of the DAG-activated pathways in cortical neurons. PKC, a main direct effector of DAG was found overactivated (**Fig. 5E**), in agreement with an excess of DAG signaling. Analysis of several other downstream effectors revealed that AKT, S6, and eIF4E were also overactivated, further supporting the alteration of DAG signaling. S6 and eIF4E are final effectors of DAG-mediated pathways, their phosphorylation levels in the brain are associated with enhanced protein synthesis rate ^66,67^. Concordantly, neuronal translation rate was found to be increased in *Dgkκ*-KO cortical neurons by puromycin incorporation assay (**Fig. 5F**). We further confirmed that DAG pathway was altered in hypothalamic primary neurons, as DGKκ is expressed at a much higher level in this neuronal type (**Fig. S5D**).

Thus, DGKκ loss phenocopies *Fmr1* KO spine/translation defects through mGluRI-DAG pathway overdrive.

### Rare missense variants in *DGKk* identified in individuals with neurodevelopmental disorders affect DGKk localization

Given the cognitive and behavioral alterations observed in the *Dgkκ*-KO mice, we investigated whether *DGKk* might contribute to neurodevelopmental disorders (NDD) in humans. To address this, we searched rare non-synonymous variants identified in NDD patients through Genematcher. We retrieved three hemizygous missense variants—p.(Thr282Ile), p.(Lys526Asn), and p.(Arg584Cys)—each detected in males with NDD who lacked other pathogenic variants in genomic analyses (exome or genome sequencing)-affecting conserved amino acids and absent from the gnomAD general population database (gnomAD v4.1.0). To assess their potential deleterious impact on protein function, we designed a functional assay. As controls, we included two presumably benign missense variants—p.(Ile612Asn) and p.(Arg851Cys)—reported in 1,079 and 1,065 hemizygous males in gnomAD, respectively, and the two catalytically inactive DGKk mutant, deleted from the C1 (DAG binding domain) or C4a (catalytic domain) domains ^63^.

To determine the functional impact of DGKk variants, we assesed the expression level and localization of DGKk p.(Thr282Ile), p.(Lys526Asn), and p.(Arg584Cys) variants (**Fig. S6A**) in neuronal and non-neuronal cells. The overexpression in HeLa cells of the variants did not lead to an altered protein level compared to WT and benign DGKk variants (**Fig. S6B**). However, compared to WT and benign variants, both DGKk p.(Thr282Ile) and p.(Arg584Cys) variants failed to localize at the plasma membrane in HeLa cells, similarly to C1 and C4a deletion mutants (**Fig. S6C**). In neurons, WT and non-pathologic DGKk variants localized at the plasma membrane of cell body as well as in dendrites. Compared to WT and benign variants, p.(Thr282Ile) and p.(Arg584Cys) and DGKk deletion mutants failed to localize efficiently in dendrites, suggesting they have an altered neuronal function (**Fig. S6D**). Further, experiments will be required to assess the pathogenicity of p.(Lys526Asn) variant.

Thus, DGKk rare loss-of-function variants identified in patients with neurodevelopmental disorders impair DGKk function in both neuronal and non-neuronal cells, thereby strengthening the candidacy of DGKk as a novel contributor to neurodevelopmental disorders in humans.

## Discussion

The identification of the RNA binding properties of FMRP has been a long-standing question in the field of FXS research. In addition to its high affinity for highly structured RNA ^22,26,27^, FMRP has also been shown to have affinity with short abundant RNA sequences ^24,25,68,69^. The FMRP binding motif “EPAP” identified here is distinct from previously described motifs and consists of a tandemly repeated GARCCGGCCCCA (more than 20 repeats) dodecamer containing a high density of m^6^A modifications. To our knowledge, FMRP displays for *Dgkκ* mRNA the best reported binding affinity ^23^. Our data suggest that FMRP’s unprecedented affinity for *Dgkκ* mRNA can be driven by the sequence of the motif itself, its repetitive nature, its high density of m^6^A modifications, and its collaborative interaction with FMRP paralogs. The “EPAP” RNA motif encodes a conserved, highly repetitive sequence (∼120 amino acids), composed of ∼80% acidic residues (E, D) and prolines, located immediately downstream of a proline stretch (**Fig. 1A**). This motif resides in exon 1, which encodes N-terminal region of DGKκ with no apparent role in catalytic function ^28,63^, implying strong selective pressure to maintain its in-frame conservation. The proline stretch upstream of the EPAP repeats, well established ribosome stalling element ^46^, is necessary to induce ribosome stalling events. The EPAP repeats are necessary to recruit FMRP in a m^6^A dependent manner in order to increase translation efficiency of DGKκ in neurons (**Fig. 2B,E,F**; **Fig. S6D**). Our findings highlight a nuanced role of FMRP that extends beyond its canonical function as a translational repressor, consistent with both a study in *Drosophila* showing that FMRP enhances translation of large proteins ^70^ and studies showing that FMRP associates with sequences where ribosomes pause — enriched in glutamate and aspartate— without directly establishing the stalling ^71,72^. Further analysis of our CLIP-seq data revealed that FMRP binding events occur on sequences encoding repeated amino acids known to promote ribosome stalling (**Fig. S7A, B**). The EPAP motif is among the motifs with highest proportion of such amino acids and the one with the greatest number of m6A consensus sites (**Fig. S7C**).

An activation of translation induced by the combined action of two factors, FMRP and m⁶A, both well described as translation repressors ^7,53,54,56,57^, seems paradoxical. Yet, ribosome pausing induced by m^6^A in coding sequence has been shown to be beneficial for the translation of certain transcripts ^52^. A similar mechanism may be at play in the stall-prone Pro-EPAP sequence, where the FMRP/m6A-mediated slowdown of translation may be beneficial to locally mitigating ribosomal collisions, ultimately enhancing productive translation (**Fig. S7D**). In this framework, FMRP emerges not simply as a brake on protein synthesis, but as a regulator of translational flow, fine-tuning ribosome behavior to modulate the expression of neurodevelopmental effectors in response to the intrinsic properties of the coding sequence. Further studies will be required to determine how FMRP regulates ribosomal collisions, either in opposition to or in concert with quality control factors such as ZNF598 or ASCC3 that intervene to rescue ribosome collision events ^73–75^ and recent work showed that this could be exploited to ameliorate FXS *Fmr1*-KO phenotypes ^76^. Moreover, it will be mechanistically insightful to define the mode of action of FMRP through structural studies of FMRP in complex with the ribosome bound to the EPAP motif, with or without m^6^A modification.

We identified differential DGKκ level alterations across the *Fmr1*-KO brain, revealing potential compensation by the FMRP paralog FXR1P, which exhibits mirror-inversed expression relative to FMRP across brain regions. This reciprocal pattern suggests functional overlap among FXR proteins and correlation of *Fmr1*-KO phenotypes with regionally specific DGKκ reductions (∼80% in the cortex and ∼50% in the hypothalamus).

Despite a low and restricted expression in certain brain areas, the loss of DGKκ revealed both neuronal and non-neuronal phenotypes shared with the *Fmr1*-KO model (**Table 1**). At behavioral level, the *Dgkκ*-KO model exhibits significant neurodevelopmental impairments, notably marked hyperactivity, reduced anxiety, hypersensitivity, and compulsive behavior, which are recurrent phenotypes of the *Fmr1*-KO mice ^77^. Consistently with other DGK mouse models ^78–80^, these findings confirm the importance of DAG-associated lipid signaling in neurodevelopment. The identification of two rare missense *DGKk* variants in males with neurodevelopmental phenotype, which disrupt DGKκ protein neuronal localization (**Fig. S6D**), positions *DGKκ* as a novel NDD candidate. Additional patients harboring deleterious variants will be required to confirm its role in NDD.

**Table 1.**
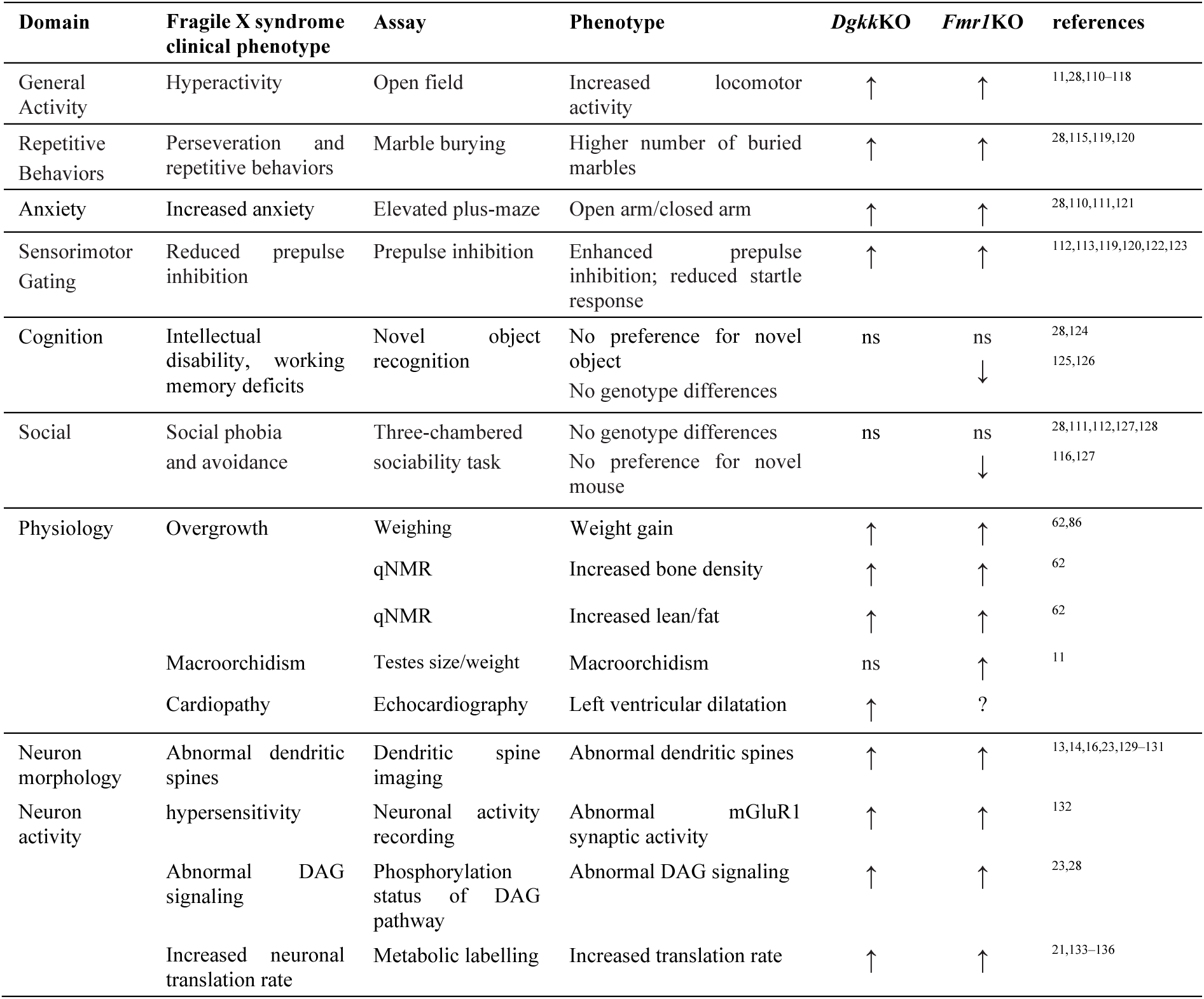
Summary of behavioral, cognitive, physiological, and cellular phenotypes of ***Dgkk*-KO vs *Fmr1*-KO knockout mice (↓ decrease; ↑ increase;** ns **no**t significant ;unknown) Metabolic labelling Increased translation rate **_↑↑_** ^21,133–136^

*Dgkκ*-KO analysis revealed the functional importance of the DGKκ enzyme in regulating neuronal DAG-signaling — a mechanism central to the adaptation to external stimuli in many cell types ^81–84—^ and how FMRP contributes to its regulation. The balance of DAG and PA plays a pivotal role in intracellular signaling ^82^ and its dysregulation, as a consequence of DGKκ lack of expression, has been postulated as an underlying pathomechanism of FXS ^23,30^. *Dgkκ*-KO neurons exhibit exacerbated DAG-signaling pathways associated with a global increase of neuronal protein synthesis rate. Increased neuronal translation rate is considered a hallmark of FXS condition and is proposed to underly LTP/LTD defects along with abnormal dendritic spine morphology leading to synaptic defects and some behavioral alterations ^12^. The exact cause of neuronal translation increase in FXS was unclear and hypothesized to be a consequence of the loss of repression by FMRP acting individually on its multiple targets ^12^. Our data unambigously demonstrate that the loss of DGKκ alone leads to such an effect. The loss of DGKκ leads also to abnormal dendritic spine morphology and an impaired mGlurI-dependent neuronal activity, similarly to the *Fmr1*-KO model. These data suggest that DGKκ loss by itself contributes to a large body of neuronal defects of FXS condition.

Beside the central nervous system phenotypes, several physiological alterations have been reported in *Fmr1*-KO mice, including lipids and glucose metabolism defects associated with increased non-fat body mass, increased muscle mass and global skeleton length, evocative of an overgrowth phenotype ^62,85^. These features are present in a proportion of FXS patients ^62,86^. FMRP was proposed to impact the hormonal system and notably hypothalamic function, which could explain the non-neuronal phenotypes in FXS ^86–90^.

To date, the molecular origin of this phenotype remains unknown. Remarkably, *Dgkκ*-KO mice mirror—and even exaggerate— the overgrowth phenotype observed in *Fmr1*-KO mice. DGKκ’s peak hypothalamic expression, coupled with its ∼50% reduction in *Fmr1*-KO mice, makes DGKκ a relevant candidate mediator of the hypothalamic phenotypes. In this context, Snord116 snoRNA, whose loss causes Prader–Willi syndrome (PWS) ^91–93^, directly targets *Dgkκ* mRNA ^94^, and its absence in PWS is associated with DGKκ upregulation ^95^. Notably, many PWS phenotypes arise from hypothalamic dysfunction, including hyperphagia, delayed puberty, and short stature, some of which overlap with FXS features ^87^. These observations, together with the hypothalamic expression of DGKκ, position it at the crossroads of both disorders, potentially contributing to their contrasting phenotypic traits.

Together, these findings position DGKκ as a key translational effector of FMRP dysfunction in FXS, opening avenues for targeted therapeutic strategies, such as DGKκ-based AAV approaches or pharmacological agents that modulate DAG/PA levels.

## Material and methods

### Animal model

*Dgkκ*^em1(IMPC)Ics^ (*Dgkκ*-KO) in C57BL/6NCrl background was obtained through Infrafrontier-I3 program by CRISPR/Cas9 deletion of exon 8 (GRCm38.p4 6897240/6897601, details available on www.mousephenotype.org. B6.129P2-Fmr1tm1.2Cidz/J (*Fmr1*-KO) in C57BL/6J was kindly given by Dr Rob Willemsen ^61^. Males from the same genotype (thereafter called *Dgkκ*-KO, *Dgkκ*-WT, *Fmr1*-KO, *Fmr1*-WT) were grouped by 3 or 4 individuals at weaning age (4 weeks), in individually ventilated cages (GM500, Tecniplast, UK), with poplar shaving bedding (Lignocell Select, JRS, Germany), and maintained under standard conditions on a 12-h light/dark cycle (7 h/19 h) with standard diet food (standard diet D04, Scientific Animal Food and Engineering, France) and water available ad libitum. Mice from the same cage received the same treatment and were transferred in the phenotyping area the next week. Animal work involved in this study was conducted according to ARRIVE guidelines and received authorization from relevant national committee (Comité National de Reflexion Ethique en Experimentation Animale) with APAFIS #17544-2018111516571205 v7 at the ICS mouse facility (Illkirch, France). Every procedure on mice was performed to ensure that discomfort, distress, pain, and injury would be minimal.

### Genotyping

Genotyping of *Dgkκ*-KO model was performed after birth on tail genomic DNA by PCR using primers CCTTCAGCAGCCTACCAACATTG and CTTCCGAAGGAACATTCCTTGG producing a 334 amplicon in WT and no product in KO, primers CCAAGGAATGTTCCTTCGGAAG and GAACATGCCCATACTCATGTACCC producing a 288 bp amplicon in WT and no product in KO, and primers CCTTCAGCAGCCTACCAACATTG and GAACATGCCCATACTCATGTACCC producing a 241 bp amplicon in KO and no product in WT. Genotyping of *Fmr1*-KO model was performed as in ^28^.

### Behavioral experiments

Behavioral experiments were conducted at 10 weeks. The behavioral experiments were performed under experimenter-independent conditions or by experimenters naive to the treatment conditions. Individual cohorts of male littermate animals (issued from *Dgkκ*^y/+^ x *Dgkκ*^+/-^ mating) were composed of WT and *Dgkκ*-KO subgroups.

### Elevated plus maze

The apparatus used is automated and made of PVC (Imetronic, Pessac, France). Two open arms (30 Å∼ 5 cm) opposite one to the other and crossed by two closed arms (30 Å∼ 5 Å∼ 15 cm). The maze has infrared captors allowing the detection of mice in the closed arms and different areas of the open arms. The mice were tested during 5 min. The number of entries and time spent in the open arms were measured to calculate an index of anxiety.

### Nest building

On the day of test, mice were singly transferred in a standard cage for the duration of nest building measurement. A block of nesting material (5 Å∼ 5 cm hemp square, Happi Mats, Utopia) was placed in the cage. Pictures were taken and visual scoring occurred at 2, 5, 24 h without disturbing the animals. The room temperature was noted when the nest was scored since nest building has a thermoregulatory function and therefore may be influenced by ambient temperatures. We used a 0-5 scale described by ^96^: 0 = undisturbed nesting material; 1 = disturbed nesting material but no nest site; 2 = a flat nest without walls; 3 = a cup nest with a wall less than the height of a dome that would cover a mouse; 4 = an incomplete dome with a wall of the height of a dome; 5 = a complete dome with walls taller than the height of a dome, which may or may not fully enclose the nest.

### Marble burying

Twenty glass marbles were placed equidistant from each other on the litter of the cages. The mice were left in contact with the marbles for a total period of 30 min. The test was terminated by removing the mice from the cages. The number of marbles was counted after 30 min. A marble was considered buried when two-thirds or more of it was covered with burying substrate.

### Novel object recognition task

Mice were tested in a circular arena (50 cm diameter and 30 cm height basin). The locomotor activity was recorded with the EthoVision XT video tracking system (Noldus, Wageningen, Netherlands). The arena was homogeneously illuminated (40 Lux) divided into central and peripheral regions. First, animals were habituated to the arena for 15 min. Each mouse was placed in the periphery of the arena and allowed to explore the apparatus. Experimenter were out of the sight of the tested animal. During the test session, the distance traveled and time spent in the central and peripheral regions were recorded. The percentage of time spent in center area was used as index of anxiety. The following day, mice were tested for object recognition in the same arena. Mice were tested for a 10-min acquisition during which they were placed in the arena in presence of a sample objects (2.5 cm diameter marble or 2 cm edge plastic dice). The time took by the animal to explore the samples was manually recorded. The following day, a 10-min retention trial was conducted. During this retention trial, the samples A and another object B (marble or dice depending on acquisition) were placed in the open field. The times took to explore the object A and object B by the animal were recorded. A recognition index was calculated as (tB / [tA + tB]) x100.

### Social recognition test

Social recognition test evaluates the preference of a mouse for a congener as compared to an object placed in an opposite compartment. This test is also used for evaluation of social memory by measuring exploration of a novel congener as compared to a familiar one. The apparatus is a transparent cage composed with a central starting compartment and two side compartments where circular grid cup (goal box) is placed at each extremity, and where the congener can be placed during testing. Testing was performed for two consecutive days. On the first day, the mouse was placed in central box then allowed to explore freely the apparatus for 10 min to attenuate their emotionality. On the second day, a C57BL/6NCrl congener from the same sex was placed in one goal box and an object was placed in the opposite one. The mouse was then placed in the starting central compartment and allowed to explore freely the apparatus for 10 min. The position of the congener and object boxes was counterbalanced to avoid any potential spatial preference. The duration of exploration of each goal box (when the mouse is sniffing the grid delimiting the goal box) was manually measured and the percentage of time the mouse took to explore the congener was used as index of social preference (recognition preference). A 10-min retention trial was then performed during which the object was replaced by a novel congener. The duration of exploration of each goal box was manually measured and the percentage of time the mouse takes to explore the congener was used as index of social memory. The social preference index (SR) is defined as (time Congener / [time Object + time Congener])x100; and the social memory index as (time novel Congener / [familiar congener + time novel Congener])x100.

### PRM-based quantification of DGKκ in tissues

Skyline software was used for the *in-silico* identification of signature peptides based on the following criteria: peptide length of 7 to 25 amino acids, excluding the first 25 N-terminal residues that could potentially constitute a signal peptide; peptides must be unique to the protein of interest (proteotypic); avoid peptides containing Cys (tends to form disulfide bonds), Met (prone to oxidation), Arg-Pro/Lys-Pro (promote missed cleavages), Asn-X-Thr/Asn-X-Ser (sites of N-glycosylation). It is important to select target peptides that are not affected by uncontrolled modifications since these lead to changes in the molecular weight of the peptide, which would appear in two forms and could no longer be considered a signature peptide. A list of mouse DGKκ peptides that meet these requirements was generated and spectral matches were found in the predictive library Prosit. Prosit was built using a deep learning algorithm that was trained with mass spectra obtained from MS analysis of synthetic reference peptides ^97^. However, experimental spectra for the signature DGKκ peptides were not found in proteomics databases (NIST, GPM, PeptideAtlas), so they needed to be validated empirically. We designed and identified on Skyline software two theoretical DGKκ -specific peptides (P06 and P07). This strategy required validation of the selected peptides and prior information such as elution time, m/z value, charge state, and fragmentation conditions, which is obtained during preliminary measurements by a global acquisition method (DDA) of the target protein produced in an overexpression system ^98^. The target peptides are then synthesized chemically with heavy isotopes that incorporate 13C- and 15N-labeled C-terminal arginine (R) or lysine (K), identical in sequence and physicochemical properties to the peptides of interest, that are needed for the actual spiking and quantification of the samples. To improve the precision of absolute abundance measurements, an external calibration curve is produced by measuring the signal intensity of the reference peptides at multiple concentration points. Spike-in of samples with heavy-labeled peptides is used as internal standards.

Frozen cell pellets or tissues were lysed in RIPA lysis buffer [50 mM Tris-HCl pH 7.4, 0.08 mM CaCl2, 100 mM KCl, 1 mM MgCl2, 1% (v/v) NP-40, 0.5% (m/v) sodium deoxycholate, 1% (m/v) SDS, 1× complete EDTA-free protease inhibitor cocktail] and protein concentration was determined using Bradford method (Bio-Rad protein assay dye reagent, #5000006). A total of 20 μg of protein extracts were migrated for 15 minutes at 150 V on a 10 % SDS-PAGE gel. Subsequently, the gel was fixed for 30 minutes in a solution containing 10% (v/v) acetic acid and 40% (v/v) ethanol, followed by staining with PageBlue Protein Staining Solution (ThermoScientific) for 1 hour. After multiple washes with ultrapure water, a band corresponding to the expected molecular weight of DGKκ was excised. Gel strips were transferred to a 96-well microplate and de-stained by washing three times with 50 mM ammonium bicarbonate (Sigma) and 50% (v/v) acetonitrile (Biosolve) for 10 minutes at room temperature with agitation at 850 rpm, followed by a wash with 100% acetonitrile under the same conditions. The gel bands were then reduced with 10 mM DTT for 30 minutes at 56°C, followed by alkylation with 50 mM IAA for 30 minutes in the dark. Gel strips were washed again using the same procedure and dried in a SpeedVac for 15 minutes.

Digestion was performed overnight at 37°C using trypsin at a concentration of 10 ng/μl diluted in 10% (v/v) acetonitrile, 50 mM ammonium bicarbonate, and 2 mM CaCl2. Peptides were extracted first with 0.2% (v/v) formic acid and 50% (v/v) acetonitrile for 15 minutes at room temperature with agitation at 850 rpm, followed by extraction with 0.2% (v/v) formic acid and 90% (v/v) acetonitrile using the same conditions, pooling two gel bands per condition. Finally, the peptide mixture was dried in a SpeedVac for approximately 3 hours, resuspended in 20 μl of 0.1% (v/v) trifluoroacetic acid (TFA), and 0.5 μl was injected into the mass spectrometer.

### Neuronal primary culture

Cortices, hippocampi, or hypothalami from *Fmr1*-WT, *Fmr1*-KO, *Dgkκ*-WT, or *Dgkκ*-KO E17.5 mouse embryos were dissected in dissection buffer containing 1xPBS, 2.56 mg/mL D-glucose, 3 mg/mL BSA, and 1.16 mM MgSO4. Each brain tissue was then incubated for 25 minutes in 100 µL of dissociation buffer composed of Neurobasal medium (GIBCO), 20 U/mL papain (Worthington), and 1 mg/mL DNase I. After centrifugation at 1000 × g for 5 minutes, the dissociation solution was discarded, and 500 µL of ovomucoid (Worthington) inhibitory solution was added and incubated for 6-7 minutes. Following this incubation, the ovomucoid solution was removed, and the cells were gently mechanically dissociated by pipetting 15 times to achieve a single-cell suspension. The dissociated cells were plated on poly-L-lysine hydrobromide-coated (P9155, Sigma) culture plates and cultured for 8 to 21 days in Neurobasal Medium supplemented with B27, penicillin/streptomycin, and 0.5 μM L-glutamine. Half of the media were changed every 2-3 days. For experiments requiring puromycin treatment, cultures were treated with puromycin solution at specified concentrations and durations as indicated.

### Western-blot

The cell lysates in the Laemmli buffer blue were heated for 5 minutes at 95°C, and 8 µL (equivalent to 15 µg of total protein) were loaded onto an SDS-PAGE denaturing gel composed of a stacking gel [4% (w/v) acrylamide/bisacrylamide 29:1 (v/v); 0.1 M Tris-HCl pH 6.8; 0.1% (w/v) SDS; 0.1% (w/v) APS; 0.01% (w/v) TEMED] and a resolving gel [10% (w/v) acrylamide/bisacrylamide 29:1 (v/v); 0.375 M Tris-HCl pH 8.8; 0.1% (w/v) SDS; 0.1% (w/v) APS; 0.01% (w/v) TEMED]. A molecular weight marker (PageRuler™ Plus Prestained Protein Ladder) was also loaded. The gel was run at 150 V for 1 hour in Tris/Glycine/SDS running buffer [0.001 M Tris-HCl; 0.2 M Glycine; 0.1% (w/v) SDS]. The transfer was performed onto a PVDF membrane (0.22 µM, Millipore) activated for 5 minutes in 100% ethanol, for 1.5 hours at 200 mA in Tris/Glycine/SDS/Ethanol transfer buffer [0.001 M Tris-HCl; 0.2 M Glycine; 0.1% (w/v) SDS; 20% (v/v) ethanol]. The PVDF membrane was then blocked with a solution of TBS (50 mM Tris-HCl pH 7.6; 150 mM NaCl) and 0.1% (v/v) Tween-20 with 5% (w/v) skim milk powder. The primary antibody diluted in TBS-0.1% (v/v) Tween-20 solution with 5% (m/v) of skim milk powder was incubated with the membrane at 4°C for 12 hours. After 3 washes of 15 minutes with gentle agitation in TBS-0.1% (v/v) Tween-20 solution, the membrane was incubated with HRP-conjugated secondary antibody (horseradish peroxidase – goat anti-mouse or rabbit J. Immunoresearch SA, 1:10000) in the same solution at room temperature for 1 hour. After 3 more washes of 15 minutes as previously described, the proteins of interest were detected using Immobilon reagent (Millipore) and visualized with a LAS600 imager (Amersham).

### qRT-PCR

Total RNA was prepared from neurons or tissue using 700 µL of TRIZOL reagent (Invitrogen) according to the manufacturer’s instructions. The RNA was then precipitated for 12 hours with two volumes of 100% isopropanol at -20°C. After centrifugation for 10 minutes at 13,000 rpm at 4°C, the supernatant was removed, and the RNA pellet was washed three times with 70% ethanol (v/v). The pellet was then resuspended in 10 µL of mQ water, and the RNA concentration was measured using a Nanodrop device. A DNase Turbo (ThermoFisher, 65001) treatment was performed on 4 µg of RNA for 30 minutes at 37°C with 1 U of the enzyme to eliminate any contamination by genomic DNA.

Reverse transcription was performed with SuperScript IV reverse transcriptase (Invitrogen) using 500 ng of purified total RNA according to the manufacturer’s instructions. qPCR was performed with the LightCycler 480 SYBR Green I Master (Roche) according to the manufacturer’s instructions using the Roche LightCycler 480 device with an activation cycle of 15 minutes at 95°C and 45 amplification cycles (15 seconds at 95°C, 30 seconds at 60°C, 30 seconds at 70°C).

### Immunofluorescence

Cells on coverslips were washed twice with 1x PBS to remove any residual media or debris. Subsequently, they were fixed with 4% (v/v) formaldehyde and 4% (m/v) sucrose in 1x PBS for 20 minutes. After fixation, the samples were washed twice again with 1x PBS. Permeabilization was carried out using 0.2% (v/v) Triton X-100 in 1x PBS for 10 minutes. For the blocking of unspecific sites, samples were incubated in a blocking buffer containing 1x PBS, 3% (m/v) bovine serum albumin (BSA), and 0.02% (v/v) Triton X-100 for 1 hour. Primary antibody incubation was performed overnight at 4°C. The primary antibody was diluted in a buffer containing 0.02% (v/v) Triton X-100 and 0.2% BSA (m/v). The next day, samples were washed three times for 10 minutes each with PBS containing 0.02% (v/v) Triton X-100 and 0.2% (m/v) BSA. Coverslips were then incubated for 1-2 hours at room temperature with fluorophore-conjugated secondary antibodies, diluted 1:1000 in the same buffer used for the primary antibody. After the secondary antibody incubation, samples were washed again three times for 10 minutes each with PBS containing 0.02% (v/v) Triton X-100 and 0.2% (m/v) BSA. Finally, the samples were mounted with Fluoromount and sealed with nail polish to prevent drying and preserve fluorescence.

### Dendritic spine morphology classification

Hippocampal primary neurons were magnetofected using NeuroMag (OzBiosciences) with a GFP plasmid at 3DIV. At 21DIV, neurons were immunolabeled using a GFP-specific antibody to enhance the signal. Selected isolated dendrites were imaged using confocal microscopy (Leica SP8X) with a 63X objective and a resolution of 4096x4096 pixels, with a step size of 0.2 µm to generate a stack of images for subsequent automated analysis using Imaris software. The following parameters were used to automatically classify dendritic spine according to their morphology: stubby: spine length < 1 µm; mushroom: spine length < 3 µm and max head width > mean neck width x 1.5; long thin: mean head width ≥ mean neck width and spine length > 1.5 µm; filopodia: spine length > 2 µm.

### MEA recording

High-density MEA Accura (3Brain) chips having 4096 electrodes were sterilized by filling the reservoir with 70% (v/v) ethanol for 20 minutes. Subsequently, the chips were rinsed three times with sterile mQ water. Once dried, the central areas of the chip, which contain the electrodes, were double-coated overnight with poly-D-lysine (P9155, Sigma) and bovine fibronectin (F1141, Sigma). Following three additional washes with mQ water, exactly 80,000 dissociated primary neuronal progenitors were seeded onto the dry chips. After allowing one hour for cell attachment, the reservoir was filled with Neurobasal Medium supplemented with B27. Throughout the experiment, the chips were maintained at 37°C with 5% CO2 in a 10 cm Petri dish with a smaller dish containing water to prevent media evaporation. Neurons were fed every two days by replacing half of the media with pre-heated media. At 21DIV, the HD-MEA chips were connected to the Biocam X device for recording neuronal activity using BrainWave 5 software. For DHPG treatment, a solution of 200 µM DHPG diluted in Neurobasal Medium supplemented with B27 was prepared and pre-heated at 37°C. For neuronal activation, while recording, half of the media in the HD-MEA chip was replaced with the DHPG-containing Neurobasal Medium supplemented with B27 to achieve a final concentration of 100 µM DHPG.

### CLIP

CLIP analysis was performed as described in (Tabet et al. 2016). Briefly, neuronal cultures at 8DIV were rinsed once with cold 1x PBS, and then proceeded for UV crosslinking (254 nm, 400 J/cm^2^, Stratalinker) on ice. Neurons from 2 wells of 6-well plate equivalent to one embryo were lysed in 400 µL of lysis buffer [50 mM Tris·HCl, pH 7.4, 100 mM KCl, 1 mM MgCl2, 0.1 mM CaCl2, 0.1% SDS, 1% Nonidet P-40, 0.5% sodium deoxycholate, 30 U of RNasin (Promega), and 10 U of DNase Turbo (Invitrogen)] and incubated for 5 min at 37 °C. Lysates were spun down at 18,000 g at 4 °C, and supernatants were precleared by incubation with 25 μL of free Dynabeads protein G (ThermoFisher) and then on 25 μL of Dynabeads protein G coupled to rabbit anti-mouse IgGs (2.5 μg) for 1 h each. The lysates were then incubated overnight with agitation on 25 μL of Dynabeads protein G coupled to 2.5 μg of anti-FMRP antibody (ab264380, Abcam). Importantly, although RNase treatment of immunoprecipitated RNA was not performed, partial RNA digestion occurs during the overnight incubation. After immunoprecipitation, the supernatants were saved for RNA extraction (WT and *Fmr1*-KO “inputs”), and the beads were washed three times with 400 µL of high-salt washing buffer [50 mM Tris·HCl, pH 7.4, 1 M KCl, 1 mM EDTA, 1% (v/v) Nonidet P-40, 0.5% (m/v) sodium deoxycholate, and 0.1% (v/v) SDS, 4 °C]. RNAs from WT and *Fmr1*-KO neurons were recovered by treatment of the beads with 0.4 mg of proteinase K in buffer (100 mM Tris·HCl, pH 7.4, 50 mM NaCl, and 10 mM EDTA) for 20 min at 37 °C. The anti-FMRP antibody was selected among several tested antibodies based on its ability to immunoprecipitated efficiently some mRNAs considered as validated FMRP targets (i.e., *Dlg4, Map1b*) compared with mRNAs considered as non-FMRP targets (*Rplp0*) in WT compared with *Fmr1*-KO neuron extracts as quantified by qRT-PCR.

For the modified iCLIP method (individual-nucleotide resolution UV crosslinking and immunoprecipitation) ^99^, the RNA crosslinked to FMRP was digested with proteinase K, generating small polypeptides cross-linked to RNA that can induce reverse transcription arrests (RT-stops) during cDNA synthesis, thereby allowing mapping of crosslink site. Detection of RT-stop events across the *Dgkκ* transcript was performed with 18 *Dgkκ*-specific primers spaced approximately 100 nucleotides apart, ensuring uniform reverse transcription coverage. RNA extracted from WT and *Fmr1*-KO immunoprecipitated samples, as well as input brain total RNA, underwent reverse transcription under identical conditions.

### RNA isolation and sequencing

RNA was extracted from WT or *Fmr1*-KO neuron inputs (Input WT) or from CLIP samples (CLIP WT and CLIP *Fmr1*-KO) using phenol/chloroform (vol/vol) followed by chloroform/isoamyl alcohol (24:1) extraction and ethanol precipitation in the presence of 0.3 M sodium acetate. Subsequently, total RNA was treated with DNase (Invitrogen, AM1907) and 500 ng of RNA was ribodepleted using the Mouse-Rat RiboPOOL kit (siTOOLs Biotech, catalog no. dp-K012-055) according to the manufacturer’s protocol, followed by reverse transcription using SuperScript IV (ThermoFisher, 18090200). For the HITS-CLIP strategy in hypothalamic neurons, reverse transcription was conducted with random hexamers following the manufacturer’s protocol (ThermoFisher, N8080127). For the iCLIP strategy in cortical neurons, a set of 18 *Dgkκ*-specific primers was utilized to cover the entire transcript of *Dgkκ* and optimize RNA-seq coverage.

To enhance primer hybridization to *Dgkκ* mRNA, we modified the reverse transcription protocol: after 5 minutes of RNA denaturation at 65°C, a hot start at 55°C was initiated by adding the reverse transcriptase mix directly to the RNA. Unlike random hexamers, which can initiate reverse transcription at multiple sites, this method allows reverse transcriptase to start only at the specific positions defined by the primers. This approach helps in blocking reinitiation when encountering FMRP polypeptide generated by proteinase K digestion, resulting in an enrichment of transcript stop at the same 5’ location. Generated cDNA was treated with 1 µL of RNase A (5 µg/mL) for 20 minutes at 37°C, then purified using RNA XP beads (Beckman Coulter) at a 1.8x ratio. The purified cDNA was sonicated in Covaris microtubes (AFA fiber Snap-cap). Finally, the fragmented cDNA was repurified with RNA XP beads and processed for MeDIP library preparation.

### EMSA

The FMRP full-length protein (Isoform 1) was prepared as a fusion protein with GST in a baculovirus system according to Bardoni et al. (1999). His–FMRP was eluted from NiNTA beads (Qiagen) in elution buffer (50 mM Tris–HCl pH 7.5, 1 mM MgCl2, 150 mM KCl, 1 mM EDTA, 1 mM DTT, 0.5M urea, 500 mM Imidazole). For binding assays, RNAs (5’GCCGGCCCCAGAGCCGG(m^6^A)CCCCGA(m^6^A)CCAG(m^6^A)CUCAGAGCCGGCCCCAGAGCCGG(m^6^A)CCCCGA(m^6^A)CCAG(m^6^A)CUCAGA(m^6^A)CCG-Biotin 3’) 5′-end labeling of RNA was performed with T4 polynucleotide kinase and [γ-32P] ATP. Labeled RNAs were purified on an 8% polyacrylamide gel (Acryl/Bisacrylamide 19/1, EU0061-B, Euromodex) in 8 M urea. Labeled RNAs (40,000 c.p.m.), renatured for 10 min at 40°C in binding buffer [10 mM HEPES, pH 7.4, 50 mM KCl, 1 mM DDT, 1 mM EDTA, 0.05% TritonX-100, 5% (v/v) glycerol, 1 µg/μl yeast tRNA, and 2 U/µl ribonuclease inhibitor] were incubated in 10 μl of binding buffer with 0–0.5 pmol at 25 °C for 30 min. The mixture was loaded on a 4% polyacrylamide gel (bisacrylamide:acrylamide 1:37.5, 2.5% (v/v) glycerol, 0.5× TBE) and migration was performed for 1h at 300 V at 4°C. The gel was then subjected to phosphoimaging.

### RNA-nHU

Protein-nHU analysis adapted from ^43^, was applied to RNA as follows: 8 days *in vitro* (8DIV) cortical neurons cultures from three embryos were washed once with cold 1x PBS, pelleted at 1000g for 5 min at 4°C, and snap-frozen in liquid nitrogen. The pellets were lysed in ice-cold lysis buffer [50 mM Hepes-KOH pH 7.5, 100 mM NaCl, 50 mM KCl, 4 mM MgCl2, 1% (v/v) TritonX, 0.5 mM DTT, 1× cOmplete EDTA-free protease inhibitor cocktail, 5 mM TCEP, 10% (v/v) glycerol, and 500 U/mL of RNasin]. Lysates underwent four rounds of sonication (4 × 20 s with 1-s pulses) on ice and were then incubated rotating at 4°C for 30 min. After centrifugation at 12,000 rpm at 4°C for 20 min, the supernatant was collected for further analysis. The total protein concentration was determined using the Bradford method (Bio-Rad protein assay dye reagent, no. 5000006) with a BSA calibration curve (MP Biomedicals, no. 160069), measured on a Bio-Rad SmartSpec 3000 spectrophotometer. Lysates were diluted to a concentration of 1 mg/ml.

For saturating streptavidin resin with biotinylated RNA probes or biotin, 50 μl of streptavidin resin was mixed with biotin or RNA probes at 20 μM concentration in 6 to 6.5 volumes of resin for 60 min. Specifically, the streptavidin resin (Streptavidin Sepharose High Performance, Cytiva) was saturated with biotinylated EPAP RNA probes, with or without m^6^A modifications, at a concentration of 20 μM in 20 volumes of resin for 60 min. After saturation, the resins were washed once with 10 volumes of holdup buffer (50 mM Tris-HCL pH 7.5, 300 mM NaCl, 1 mM TCEP, 40 U/mL RNasin, 0.22-μm filtered), and any remaining streptavidin beads were blocked with 100 μM biotin in 10 volumes of holdup buffer for 10 min. Finally, the resins were washed twice with 10 volumes of holdup buffer. The nHU mixture was then incubated at 4°C for 1 hour. After incubation, the resin was separated from the supernatant by brief centrifugation (15 sec, 2000 g), and half of the supernatant was immediately pipetted off to prevent resin contamination.

### RNA pull-down

For beads coupling, 45 µL of Dynabeads MyOne Streptavidin (ThermoFischer, 65001) were incubated with RNA binding buffer [50 mM HEPES, pH 7.5, 150 mM NaCl, 0.5% (v/v) NP40, 10 mM MgCl2] containing 0.8 U/µL RNasin (Promega) for 30 minutes on ice. After removing the supernatant to eliminate RNAse-RNasin complexes, the beads were blocked in RNA binding buffer supplemented with 50 µg/mL yeast tRNA for 1 hour at 4°C under head-over-tail agitation. Following two washes with RNA binding buffer, 5 µg of RNA probes were added to the beads in 600 µL of RNA binding buffer and incubated for 1 hour at 4°C under head-over-tail agitation. Subsequently, RNA-bead complexes were washed twice with RNA binding buffer and twice with lysis buffer [50 mM Tris-HCl, pH 7.5, 150 mM KCl, 1 mM MgCl2, 1 mM EDTA, 1% (v/v) NP40, 0.5 mM DTT, 1× complete EDTA-free protease inhibitor cocktail, 2.5% (v/v) glycerol, RNasin 200 U/mL].

Pellets of cortical neurons, equivalent to 5 embryos, were lysed in 300 µL of lysis buffer and incubated for 15 minutes at 4°C under head-over-tail agitation. After centrifugation at 1000 x g for 10 minutes at 4°C, the pellets corresponding to the nuclei fraction were resuspended in 300 µL of lysis buffer, sonicated (5 x 30 seconds pulses), and combined with the cytosol fraction. Lysates were then centrifuged at 18,000 x g for 10 minutes at 4°C, and protein concentration was determined using the Bradford method. Subsequently, 500 µg of protein lysate was incubated with RNA-bead complexes in 600 µL of lysis buffer for 1 hour under head-over-tail agitation. Finally, beads were washed three times with lysis buffer and stored dried at -20°C until proteomic analysis.

### Northern-blot

Cell pellets were lysed on ice in 300 µL lysis buffer consisting of 50 mM Tris-HCl pH 7.5, 5 mM MgCl2, 150 mM KCl, 1 mM DTT, 0.5% NP-40, cycloheximide (100 µg/mL final), and protease inhibitor cocktail. After gentle pipetting, lysate was centrifuged at 7000 x g for 8 min and RNA concentration was quantified using the Qubit BR RNA assay (Thermo Fisher).

An aliquot corresponding to 25 µg total RNA was adjusted to 300 µL using lysis buffer, and RNase A digestion was initiated by addition of 0.8 µg RNase A (Thermo Fisher, R1253). Samples were incubated at 25 °C for 20 min. Digestions were stopped by addition of TRIzol (Invitrogen) and RNA was extracted according to the manufacturer’s instructions. Ten micrograms of purified RNA were loaded per lane for subsequent Northern blot analysis. After migration, gel was stained with ethidium bromide for imaging and RNA was electrotransfered onto a positively charged nylon membrane (Cytivia, 25006218) and proceeded twice to UV crosslinking (254 nm, 1200 J, Stratalinker).

PCR product (∼ 400 nt) spanning the region of interest were radioactively labeled using random priming. Briefly, 100 ng of purified DNA probe were mixed with 2 µL of random hexamers (Thermo Fisher, N8080127) in a final volume of 15 µL, heated to 95 °C for 5 min, and rapidly chilled on ice for ≥2 min. The denatured probes were then combined with 6 µL of cold dNTP mix (0.5 mM each of dATP, dGTP, and dTTP), 5 µL of 10x Klenow buffer, 5 µL of [α-32P]-dCTP, 1 µL of Klenow fragment (Thermo Fisher), and nuclease-free water to a final reaction volume of 50 µL. The labeling reaction was incubated at 37 °C for 2 h. Probes were then purified using MicroSpin G-50 columns (Cytivia, 27533001).

Following labeling, probes were diluted by adding 760 µL of nuclease-free water. For hybridization, 400 µL of probe (∼50 ng DNA) were heated to 95 °C for 5 min to denature, then immediately added to ∼5 mL of hybridization solution (6x SSC [150 mM NaCl, 15 mM sodium citrate], 0.5% SDS). Membranes were incubated with the denatured probe overnight at 60 °C. Membranes were washed twice with SSC 2x, and once with SSC 2x, 0.1 % SDS for 10 min at 60°C. Finally, the membranes were then subjected to phosphoimaging.

### Peak calling, filtration and annotation

The analyses of the CLIP-Seq dataset were implemented on Galaxy ^100^. The protocol was adapted from ^101^. In brief, adapters were removed and reads from fastq files of CLIP-Seq (IP) and bulk RNA-Seq (Input) were trimmed using Trimmomatic ^102^ (Galaxy Version 0.39+galaxy2). Reads were then aligned on the Mouse reference genome mm39 using STAR ^103^ (Galaxy Version 2.7.11b+galaxy0). Duplicates were removed with Sambamba ^104^ (Galaxy Version 1.0.1+galaxy2). Peaks were called using PEAKachu (https://github.com/tbischler/PEAKachu) (Galaxy Version 0.2.0+galaxy0), with the bam of bulk RNA-Seq data as control. Simultaneously, we estimated gene expression by counting reads aligned on each gene as define in the reference annotation Gencode v. M38 with FeatureCounts ^105^ (Galaxy Version 2.1.1+galaxy0). Only genes with a count of at least 50 reads in either CLIP-Seq data or RNA-Seq data were further analyzed. We performed a differential expression analysis between RNA-Seq and CLIP-seq transcriptomic profiles with limma (limma-voom) ^106^ (Galaxy Version 3.58.1+galaxy0).

The peaks detected with PEAKachu (https://github.com/tbischler/PEAKachu) were further filtered and annotated. We first removed peaks for which expression in the RNA-Seq was null (noted Inf by PEAKachu). For annotation, we used the BioMart tool ^107^ to extract coordinates, strand for canonical transcripts of the mouse genome annotation. We then only kept peaks overlapping annotated exons on the same strand with IntersectBed for the Bedtools suite ^108^. We further kept peaks overlapping genes found over-expressed in the CLIP-Seq dataset when compared to the RNA-Seq dataset (with limma).

To convert the RNA sequence under the peaks, we used the R package geno2proteo ^109^. We then counted the frequency each amino acid under the remaining peaks (**Supplementary Table S2**).

## Data availability

Clip-seq data are available in GEO repository under accession number GSE324369.

## Acknowledgments

We thank Andrea Accogli (McGill U.), Vincenzo Salpietro Damiano (UCL Institute of Neurology, UK), SeSong Jang (Seoul U), Patrick Edry (Lyon U) for sharing data about DGKk variants. We are deeply grateful to Prof Jean-Louis Mandel for constant support throughout this project, Marie-Christine Birling and the members of the Institut Clinique de la Souris for their invaluable assistance in generating the *Dgkκ*-KO model and performing behavioral studies. We also acknowledge Bernard Jost and Stephanie Legras for their expert guidance on sequencing strategies and sequence analysis, Erwan Grandgirard for imaging support, and Bastien Morlet for his help with PRM studies. We are particularly indebted to Nicolas Charlet-Berguerand, Frank Martin, Juliette Godin, Albert Weixlbaumer for their insightful discussions and suggestions. Our sincere thanks go to Sirine Souali-Crespo for her support in anatomical studies and to Nicolas Zerbe and the IGBMC core facilities for their technical assistance. Finally, we thank our colleagues in the TMN department for their support and interest throughout this work.

## Author contributions

OC, BZ, LN, GG designed and performed experiments and helped writing the manuscript. ND and AP performed experiments. AP helped designing the experiments and writing manuscript. TM and OC designed and performed bioinformatic analyses. HM supervised the project, designed and performed experiments and wrote the manuscript.

## Supplementary figure legends

**Figure S1:**
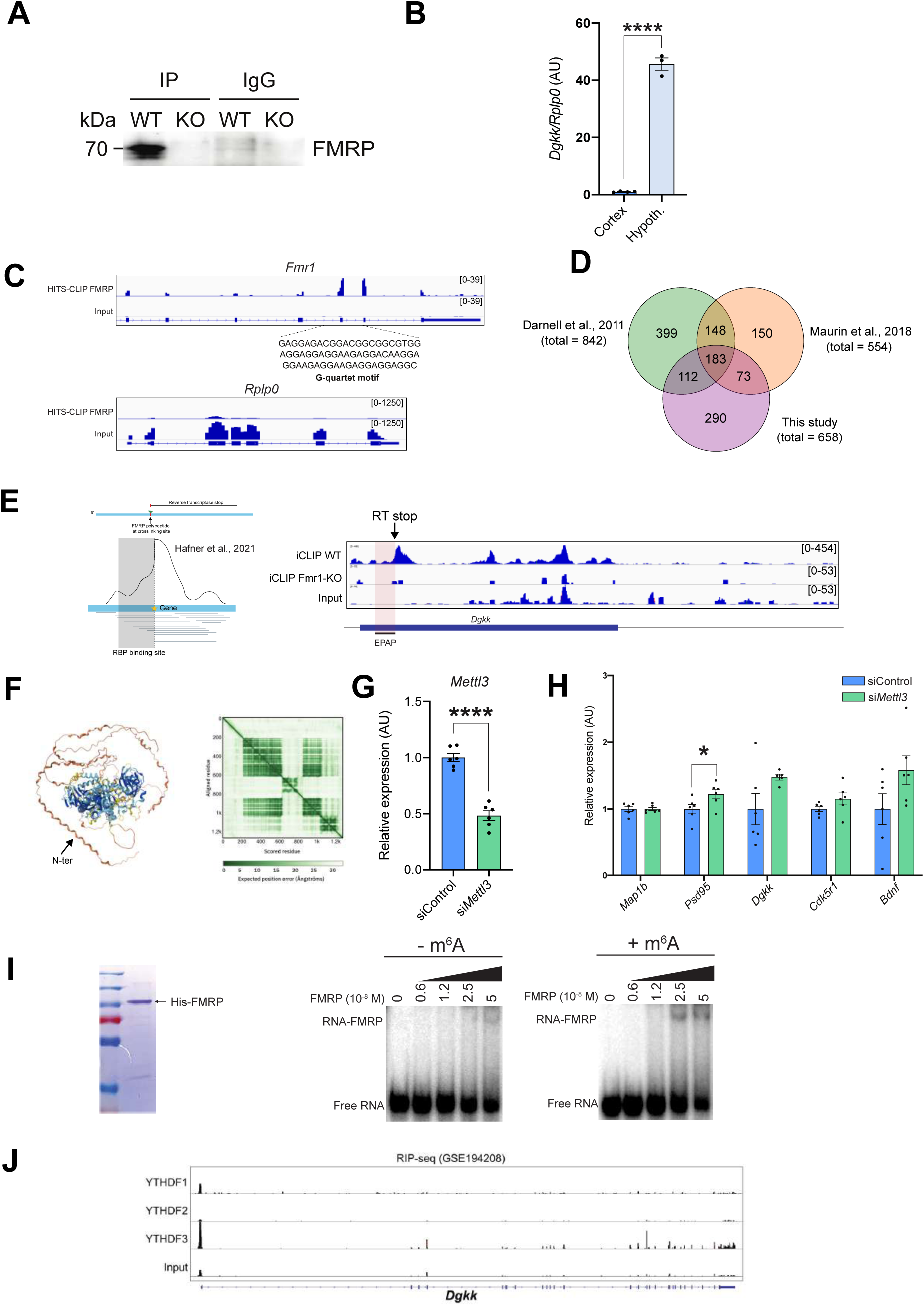
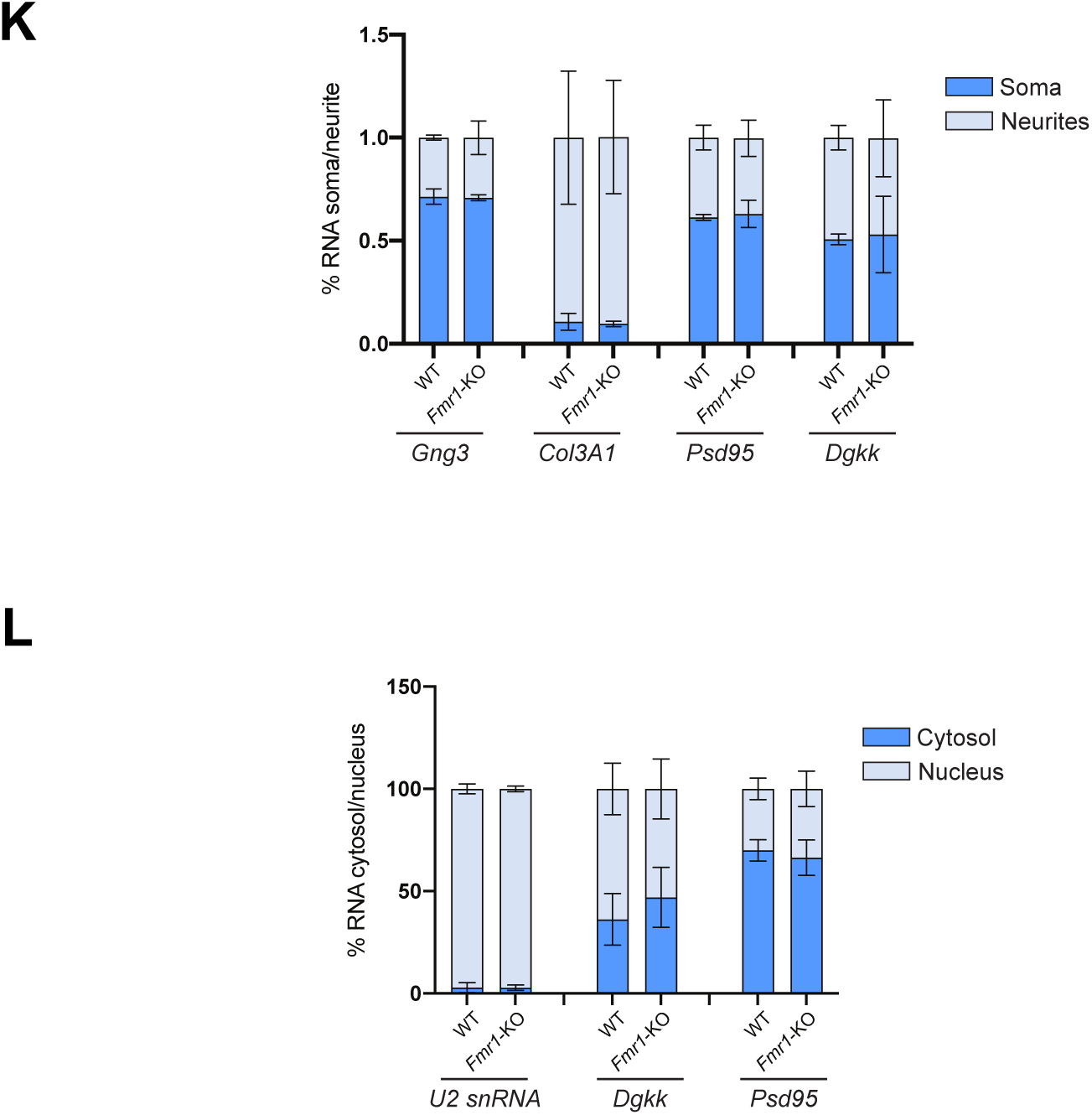
FMRP preferentially binds repetitive m⁶A-modified coding sequences and regulates DGKκ translation. A) Validation of FMRP immunoprecipitation in cortical neurons using an FMRP-specific antibody. *Fmr1*-KO neurons and IgG antibodies were used as negative controls. B) qRT–PCR analysis of *Dgkκ* expression in primary cortical and hypothalamic neurons at DIV8. Relative expression levels were calculated using the ΔΔCT method with *Rplp0* as the housekeeping gene. Data are presented as mean ± SEM and were analyzed using an unpaired Student’s t-test. ****P < 0.0001. C) IGV representation of *Fmr1* (top panel) and *Rplp0* (bottom panel) mRNAs showing HITS-CLIP-seq profiles of FMRP relative to input. D) Venn diagram illustrating the overlap between mRNA targets identified in this FMRP CLIP study and targets reported in two independent published datasets ^25,138^. Shared and study-specific targets are indicated. For the ^25^ CLIP study, the FMRP target list was restricted to mRNAs containing more than one FMRP peak and supported by at least 50 reads. E) iCLIP-seq (individual-nucleotide resolution UV crosslinking and immunoprecipitation) ^99^ allows identification of WT-specific RT-stop events downstream of EPAP that were absent in both *Fmr1*-KO and input RNA samples. IGV screenshot of FMRP iCLIP-seq data on *Dgkκ* mRNA. The red box highlights a WT-specific peak corresponding to a truncation event indicative of an FMRP binding site, as predicted based on iCLIP read-through profiles ^139^. F) AlphaFold-predicted structure of human DGKκ, highlighting the intrinsically disordered region corresponding to the EPAP domain. G) qRT-PCR analysis of *Mettl3* expression in cortical neurons treated at DIV5 for 72 h with Accell control siRNA or siRNA targeting *Mettl3*. Fold changes were calculated using the ΔΔCT method with ActB as the housekeeping gene. Data are mean ± SEM and were analyzed using multiple t-tests. ****P < 0.0001. H) qRT-PCR analysis of the indicated genes in cortical neurons treated at DIV5 for 72 h with Accell control siRNA or siRNA targeting *Mettl3*. Fold changes were calculated using the ΔΔCT method with *ActB* as the housekeeping gene. Data are mean ± SEM and analyzed using multiple t-tests comparing expression in siControl and si*Mettl3* conditions for each gene. *P<0.05. I) EMSA using purified human His-tagged FMRP isoform 1 and radiolabeled EPAP RNA probes with or without m⁶A modification. J) IGV representation of *Dgkκ* mRNA showing RIP-seq profiles of YTHDF proteins in hippocampal neurons ^42^. K) qRT-PCR analysis of cytosolic and nuclear RNA fractions to assess mRNA nuclear export of the indicated genes in WT and *Fmr1*-KO brain tissue. *U2* snRNA was used as a non-exported nuclear control. L) qRT-PCR analysis of neurite and soma fractions to assess mRNA transport in WT and *Fmr1*-KO cortical neurons. *Gng3* was used as a poorly transported control mRNA and *Col3a1* as a highly neurite-enriched control.

**Figure S2:**
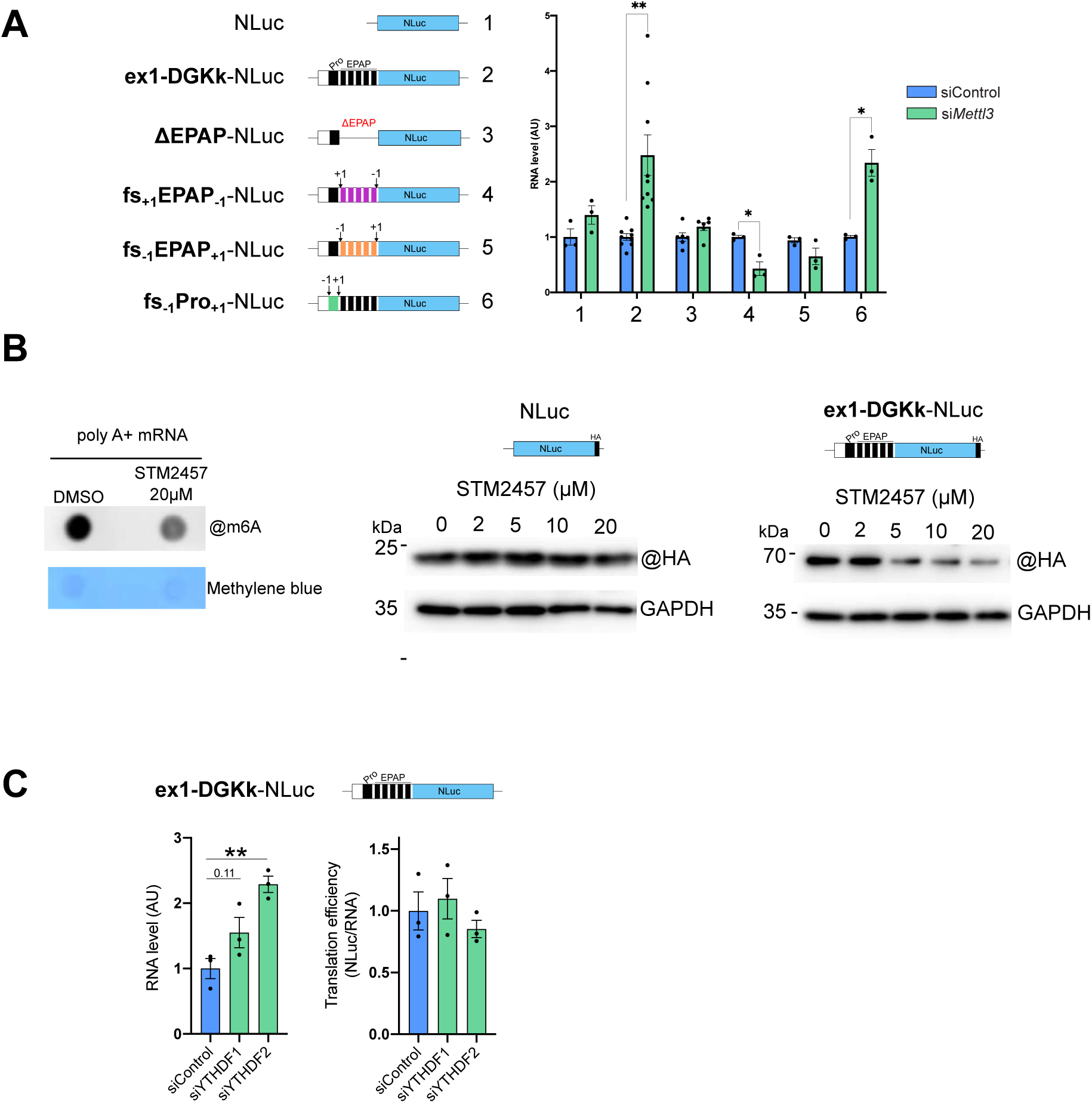
m⁶A modification in the EPAP repeats regulates DGKκ translation. A) qRT-PCR analysis in HeLa cells co-transfected for 24 h with the indicated constructs and control siRNA or siRNA targeting *METTL3*. GAPDH was used as the housekeeping gene. Data are mean ± SEM and analyzed using multiple t-tests comparing expression in siControl and si*METTL3* conditions for each gene. *P<0.05; **P<0.01. B) Dot blot analysis using an anti-m⁶A antibody on 1 µg of poly(A)+ RNA isolated from HeLa cells treated with DMSO or 20 µM STM2457 for 24 h. Methylene blue staining was used as a loading control. Representative western blots of HA-tagged constructs expressed in HeLa cells treated with increasing concentrations of STM2457 for 24 h are shown. GAPDH was used as a loading control. C) qRT-PCR analysis of Nluc mRNA in HeLa cells transfected for 24 h with control siRNA or siRNA targeting *YTHDF1* or *YTHDF2*. GAPDH was used as the housekeeping gene. Translation efficiency of Ex1-DGKκ–NLuc was measured by normalizing luciferase activity to RNA levels of the corresponding construct. Data are presented as mean ± SEM and analyzed by one-way ANOVA with comparison to the siControl condition. *P < 0.05.

**Figure S3:**
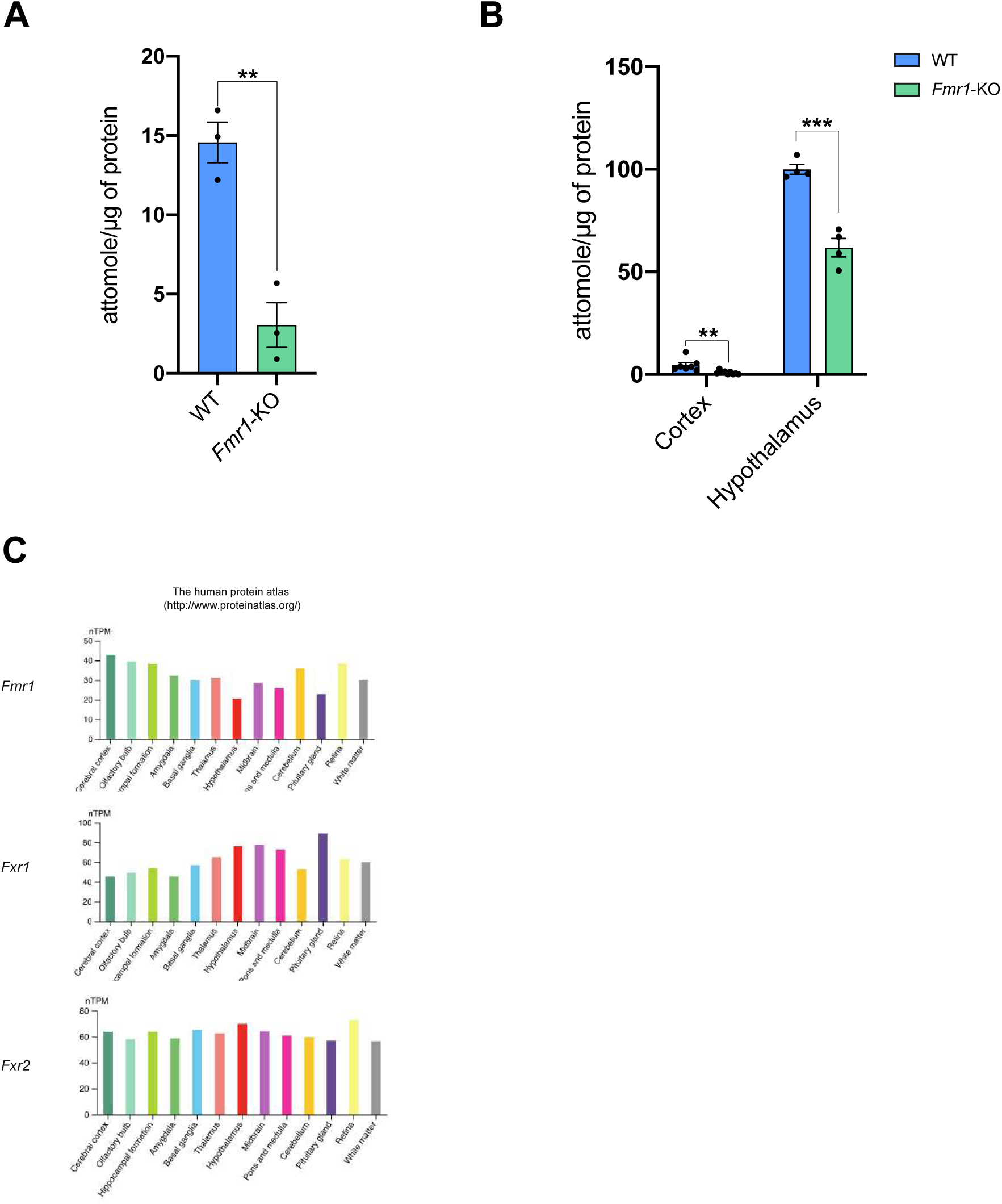
FXRs expression across the brain. A) PRM-targeted proteomic quantification of DGKκ in primary cortical neurons at DIV7 from WT and *Fmr1*-KO mice. Each dot represents an individual culture. Data are mean ± SEM and were analyzed using an unpaired Student’s t-test. **P < 0.01. B) PRM-targeted proteomic quantification of DGKκ in cortex and hypothalamus from WT and *Fmr1*-KO mice. Each dot represents an individual mouse. Data are mean ± SEM and were analyzed using an unpaired Student’s t-test. **P < 0.01, ***P<0.001. C) Expression pattern of FXR proteins in the mouse brain. Data obtained from www.proteinatlas.org.

**Figure S4:**
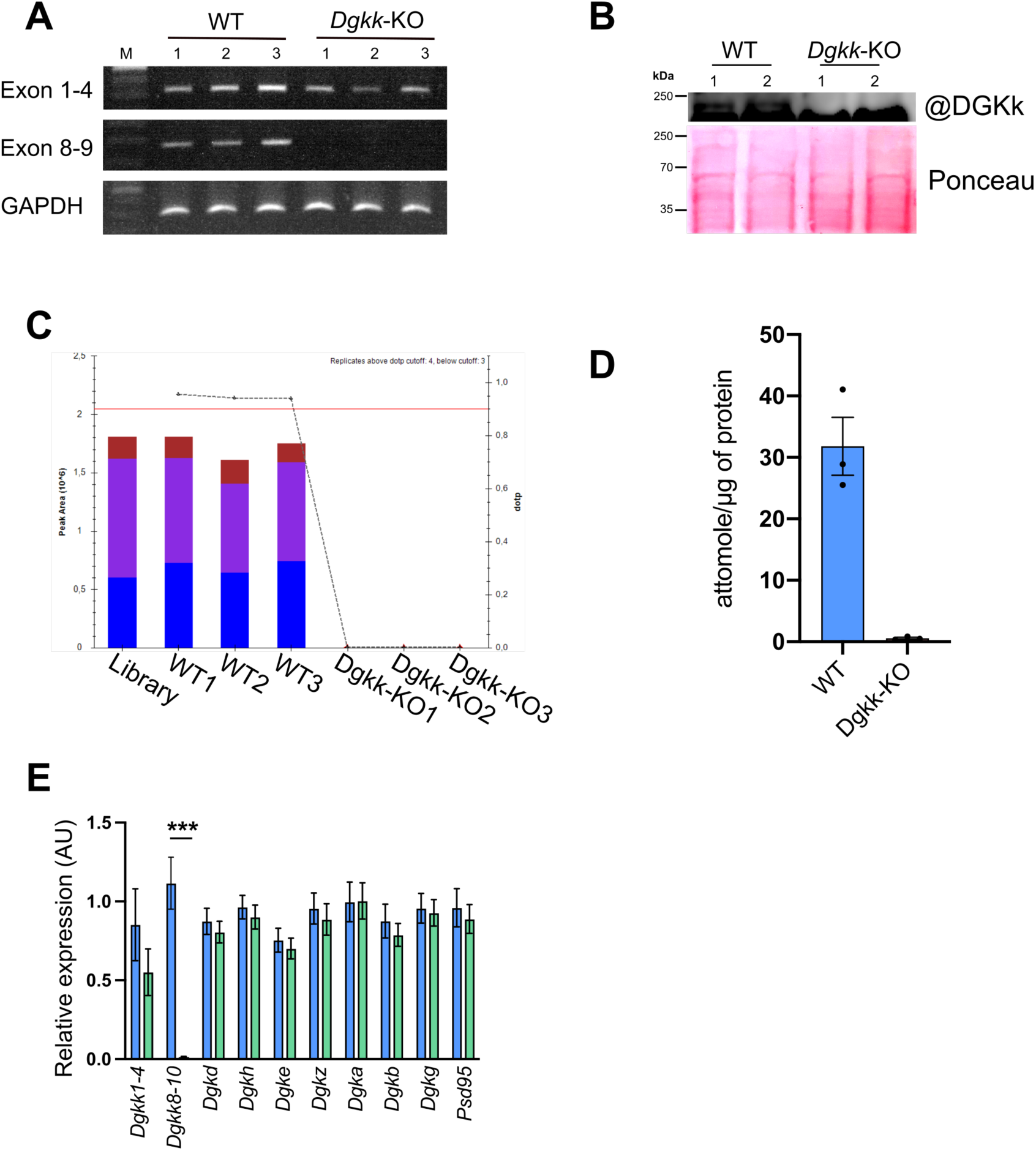
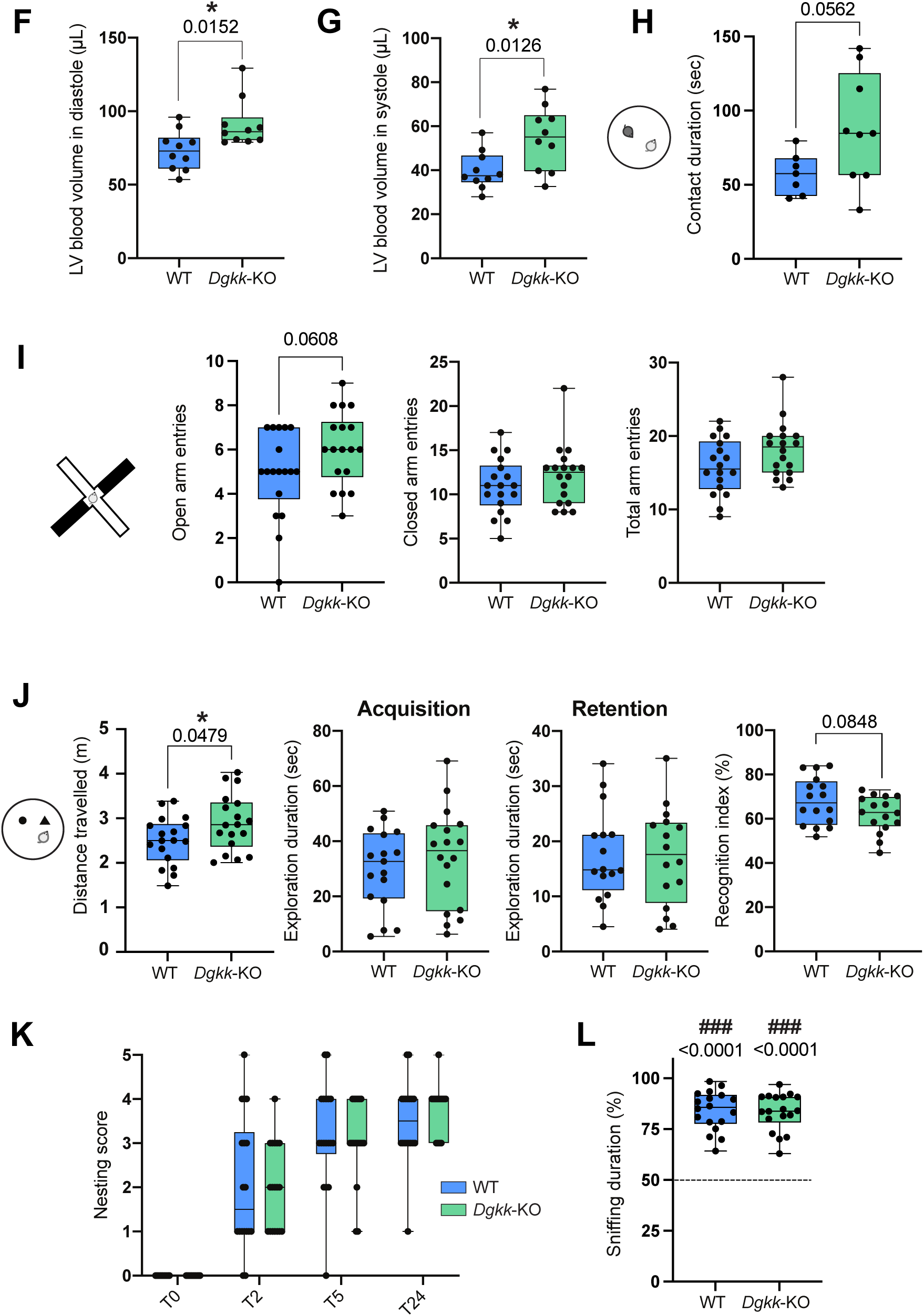
Behavioral analyses and echocardiography of *Dgkκ*-KO mice. A) RT-PCR analysis of exons 1–4 and 8–9 of the *Dgkκ* gene in WT and *Dgkκ*-KO mice. *Gapdh* RT-PCR was used as a loading control. B) Western blot of hypothalamic lysates from WT and *Dgkκ*-KO mice using a partially DGKκ-specific antibody. C–D) Detection and quantification of a DGKκ-specific peptide by targeted proteomics in hypothalamic lysates from WT and *Dgkκ*-KO mice. C, representative detection. D, Quantification of DGKκ-specific peptide levels. E) qRT-PCR analysis of 9 Dgk isoforms and *Psd95* in WT and *Dgkκ*-KO mouse brains. Data are presented as mean fold change ± SEM and analyzed using unpaired Student’s t-test. ***P<0.001. F–G) Echocardiography measurement of left ventricular blood volume during diastole (F) and systole (G). H) Social interaction test. Number of interactions between the test mouse and a novel conspecific. I) Elevated plus maze test. Measure of open arms and closed arms in a “+”-shaped maze. Total arm entries were counted to assess general locomotor activity. J) Novel object recognition test. Exploration duration during the habituation, acquisition, and retention phases. Recognition index was calculated to assess episodic memory. K) Nest-building assay. Nests were scored at 2, 5, and 24 h using a 0–5 scale as described by ^96^: 0 = undisturbed nesting material; 1 = disturbed material but no nest site; 2 = flat nest without walls; 3 = cup nest with wall < ½ the height of a dome; 4 = incomplete dome with wall ½ the height of a dome; 5 = complete dome with walls > ½ the height of a dome, which may or may not fully enclose the nest. L) Social memory (percentage of exploration of a novel vs familiar congener). F-K) Each dot represents an individual mouse. Data are mean as median with interquartile range with minimum and maximum values and were analyzed using an unpaired Student’s t-test. *P < 0.05. L) Data are expressed as median with interquartile range with minimum and maximum values and analyzed using one group t-test. ### p<0.0001 vs chance (50%).

**Figure S5:**
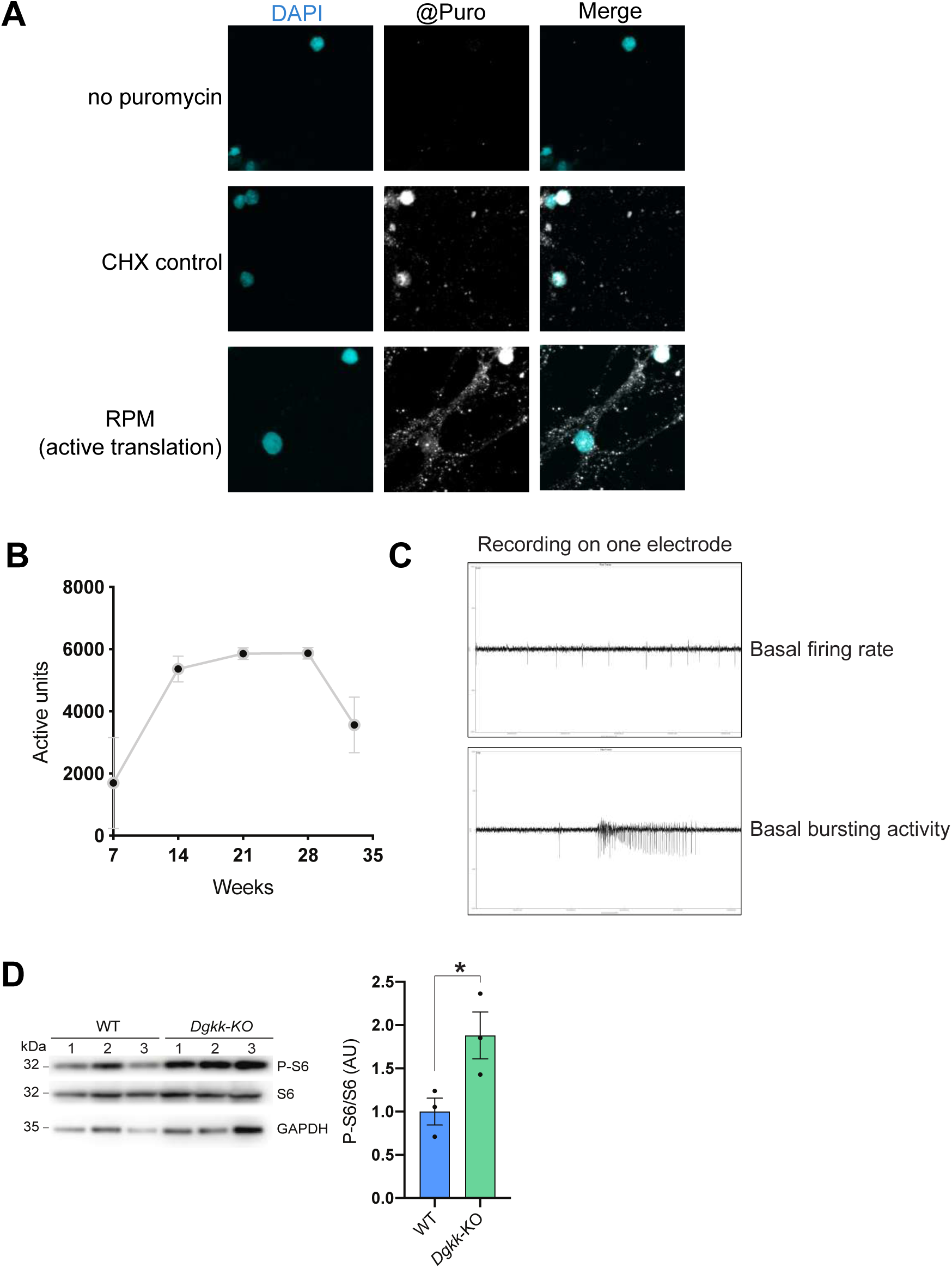
DGKκ regulates translation and neuronal electrical activity. A) Ribopuromycylation assay in hippocampal neurons. Top panel: no puromycin control. Middle panel: neurons pre-treated with 1.6 mM cycloheximide prior to ribopuromycylation to verify that puromycin signal is translation-specific. Bottom panel: ribopuromycylation signal. B) Number of active units recorded on MEA chips over time, reflecting neuronal activity from 7 DIV to 35 DIV. C) Representative raw signal showing action potentials at basal level and during spontaneous bursting activity on a single electrode. D) Western blot analysis of lysates from WT and *Dgkκ*-KO hypothalamic neurons at 7 DIV, showing total and phosphorylated S6 protein. GAPDH was used as a loading control. Quantification: phospho-S6 signal normalized to total S6 and expressed relative to WT. Each dot represents an independent culture. Data are mean ± SEM and analyzed using unpaired Student’s t-test. *P < 0.05.

**Figure S6:**
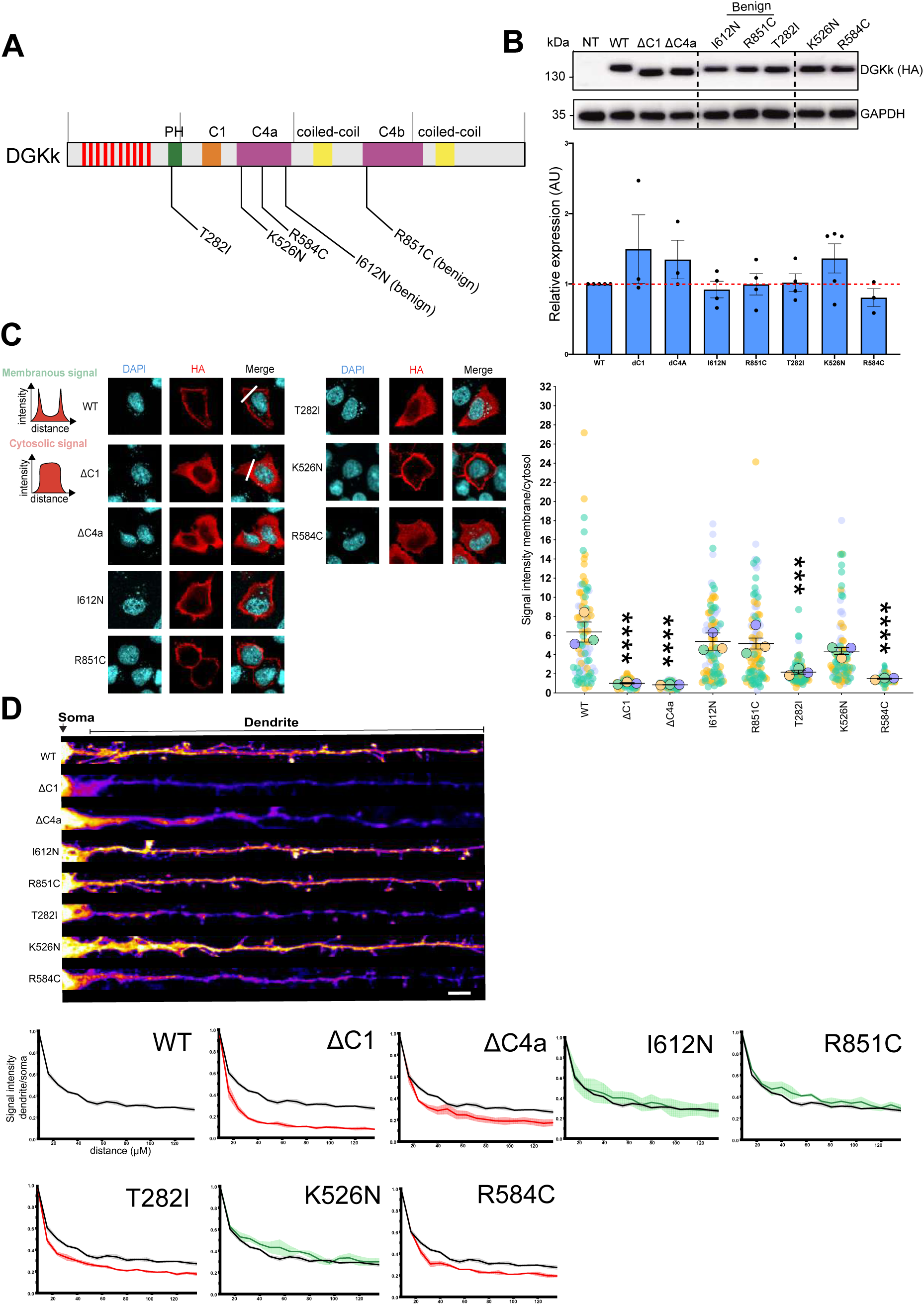
Functional impact of DGKk variants identified in individuals with NDD. A) DGKk domains and location of the identified missense variants. B) Representative western blot of overexpressed DGKk variants found in patients with ID. Quantifications of the HA-DGKk signal were normalized against GAPDH protein signal and are presented relative to the WT. C) Cellular localization of DGKk and impact of patient variants on its membrane localization. HeLa cells were transfected with the indicated constructs. DGKk catalytic domain deletion and benign mutation constructs were used respectively as a positive and negative controls. The histogram represents the signal intensity ratio membrane/cytosol obtained by measuring the signal intensity of a line passing through the cytosol (as indicated). A total of 100 cells from 3 different different cultures were counted and represented with different color symbols for each culture. Scale bar: 10 µM. D) Neuronal localization of DGKk and impact of patient variants on its dendritic localization. Neurons were magnetofected at 3DIV and fixed at 8DIV. DGKk catalytic domain deletion and benign variant constructs were used as positive and negative controls, respectively. The quantification indicates the signal intensity ratio dendrite/soma over a distance of 120 µM. A total of 12 to 16 neurons from 3-4 cultures were quantified for each condition. Scale bar: 20 µM. Data are mean ± SEM.

**Figure S7:**
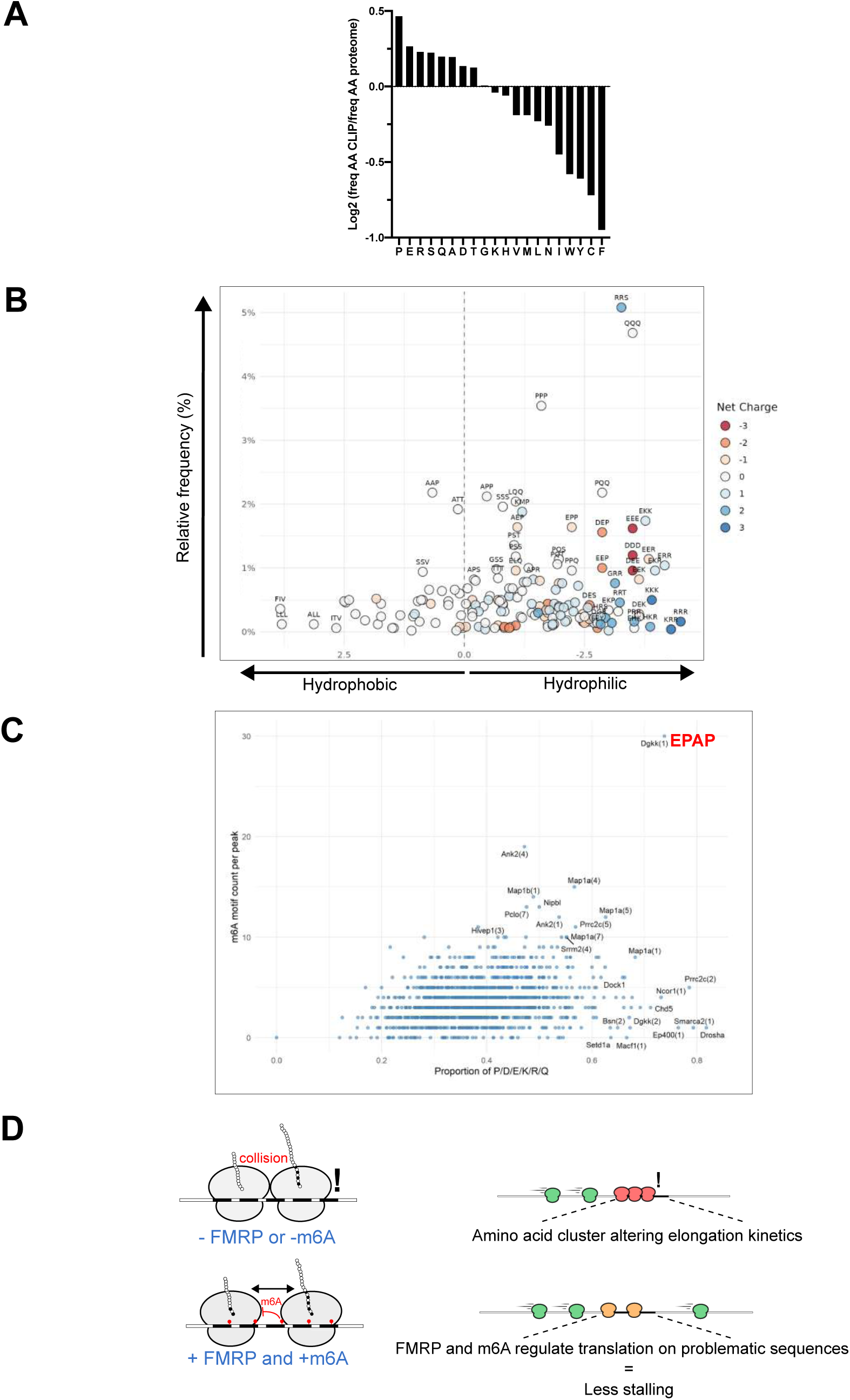
FMRP mostly binds sequences coding for stalling-prone sequences. A) Frequency of amino acids encoded by sequences enriched in FMRP HITS-CLIP-seq targets relative to their frequency in the mouse proteome. B) Hydrophobicity versus frequency of enriched tripeptides identified in FMRP-bound peptide sequences. Scatter plot showing the enrichment of unordered tripeptide motifs detected in amino-acid sequences derived from FMRP CLIP–enriched transcripts. Tripeptides were scanned in sliding windows of 30 amino acids, and motifs detected ≥2 times within the same window were collapsed into non-ordered compositions (e.g., EPE/PEE/EEP → “EEP”). The y-axis reports the fraction of enriched windows containing each motif across all FMRP-associated sequences. The x-axis shows the mean Kyte–Doolittle hydrophobicity, plotted on an inverted scale to distinguish hydrophobic (left) from hydrophilic (right) motifs; the dashed vertical line marks hydrophobicity 0. Point color indicates net charge (K/R: +1; D/E: –1), and selected motifs with high co-occurrence or extreme hydrophobicity are labelled. C) Correlation between amino acid composition and m^6^A consensus density. Each data point represents a genomic peak from the EPAP dataset. The x-axis shows the cumulative proportion of P, D, E, K, R, and Q amino acids within the protein sequence encoded by each peak, and the y-axis displays the number of m^6^A motif per peak. D) Proposed model for the coordinated action of FMRP and m⁶A on amino acid clusters altering translational elongation process.

